# Human FAM111A inhibits vaccinia virus replication by degrading viral DNA-binding protein I3 and is antagonized by poxvirus host range factor SPI-1

**DOI:** 10.1101/2023.02.04.527148

**Authors:** Junda Zhu, Xintao Gao, Zihui Zhang, Yining Deng, Shijie Xie, Shuning Ren, Yarui Li, Hua Li, Kang Niu, Shufang Fu, Yinü Li, Bernard Moss, Wenxue Wu, Chen Peng

## Abstract

Poxviruses are large double-stranded DNA viruses that infect a wide range of animals including humans. Since the eradication of smallpox, other members of the poxvirus family, such as monkeypox virus (MPXV) are still posing a great threat to public health. Vaccinia virus (VACV) is a prototypic poxvirus used as the vaccine strain for smallpox eradication. VACV encodes a serine protease inhibitor 1 (SPI-1) conserved in all orthopoxviruses, which has been recognized as a host range factor for modified vaccinia virus Ankara (MVA), an approved smallpox vaccine and a promising vaccine vector. FAM111A, a nuclear protein that regulates host DNA replication, was shown to restrict the replication of VACV-ΔSPI-1 in human cells. Nevertheless, the detailed antiviral mechanisms of FAM111A were unresolved. Here, we show that FAM111A is a potent restriction factor for VACV-ΔSPI-1 and MVA. Deletion of FAM111A rescued the replication of MVA and VACV-ΔSPI-1 and overexpression of FAM111A significantly reduced viral DNA replication and virus titers but did not affect viral early gene expression. The antiviral effect of FAM111A necessitated its trypsin-like protease domain and DNA binding domain but not the PCNA-interacting motif. We further discovered that FAM111A translocated into the cytoplasm upon VACV infection and this process was mediated by the cGAS-STING signaling pathway. Infection-triggered FAM111A degraded the nuclear pore complex via its protease activity, translocated to the cytoplasm, and interacted with and promoted the degradation of virus DNA binding protein I3 in a DNA-dependent manner. Interestingly, the protease activity of FAM111A was only needed for nuclear export but not I3 degradation as further analysis showed I3 was degraded through autophagy. Moreover, VACV SPI-1 was found primarily in the nucleus of infected cells and antagonized FAM111A by prohibiting its nuclear export. MPXV and lumpy skin disease virus SPI-1s also inhibited human FAM111A. Our findings reveal the detailed mechanism by which FAM111A functions to restrict a cytoplasmic DNA virus and provide explanations for the immune evasive function of VACV SPI-1.

## Introduction

During their co-evolution, viruses and their hosts have evolved numerous strategies to antagonize each other. Upon viral infection, cells initiate various programs, such as the induction of type I interferons and the expression of an armory of antiviral factors that can restrict viral invasion, gene expression, DNA replication, and virion assembly. More recent findings point to the fact that some host defense factors are multifunctional and can display potent antiviral activity in a way that is distinct from their regular cellular functions^1–4^. One such example is FAM111A (family with sequence similarity 111 member A), a protein found exclusively in mammals harboring a fully functional trypsin-like C-terminal protease domain^5^. FAM111A has been shown to prevent replication forks from stalling via its protease domain and disease-associated mutants display aberrant nuclear morphology due to the disruption of nuclear pore complex (NPC) as a result of its hyperactive protease activity^6, 7^. Recently, more studies have pinpointed the function of FAM111A in restricting virus infection. Fine et al. identified FAM111A as a host restriction factor for SV40 and showed FAM111A was specifically targeted by the C-terminal region of Simian Virus 40 (SV40) large T (LT) antigen^6, 8^. Moreover, Panda et al. identified FAM111A as a restriction factor for VACV-ΔSPI-1 through genome-wide RNAi screening^9^. Nevertheless, detailed mechanisms by which a nuclear protein FAM111A restricts viral infections, especially infections of poxviruses that replicate exclusively in the cytoplasm, remain largely elusive.

Poxviruses are large cytoplasmic DNA viruses, and as a family, can infect a wide variety of different animal species. The most notorious members of the poxvirus family include variola virus, the causative agent of smallpox, and MPXV, which is responsible for the recent global outbreaks of monkeypox infections since 2021^10–13^. VACV was used as the vaccine strain for the eradication of smallpox and is also the prototypic poxvirus that is best characterized^14, 15^. The genome of VACV encodes for more than 200 genes, nearly half of which are involved in host-virus interactions and a third of them responsible for evading host antiviral defenses^16^. VACV encodes for three serine protease inhibitors (SPI-1, SPI-2, and SPI-3). Deletion of SPI-1 from VACV caused a host range defect and insertion of it into MVA, the highly attenuated vaccine strain unable to replicate in most mammalian cells, was able to rescue MVA’s replicative capability in mammalian cells such as BS-C-1 and MRC-5^17, 18^. As a previous study identified FAM111A as one of the major cellular targets of SPI-1, more efforts are needed to characterize the role of SPI-1 in antagonizing FAM111A’s antiviral effect on vaccinia virus^9^.

Here, we demonstrate that FAM111A, a component of the replication fork primarily localized in the nucleus under normal conditions, translocated into the cytoplasm in VACV-infected cells upon the activation of the cGAS-STING signaling pathway by VACV infection. The export of FAM111A necessitated the disruption of NPC via its protease activity. Cytoplasmic FAM111A colocalized with viral factories, bound and mediated the degradation of a viral DNA binding protein I3 through autophagy, leading to the arrest of viral DNA replication, post-replicative protein expression, and eventually virus replication. Upon infection, SPI-1 entered the nucleus of infected cells and prevented the nuclear export of FAM111A by inhibiting the peptidase activity of the latter, and a functional SPI-1 reactive-site loop (RSL) was indispensable for this inhibitory effect^19^. Our findings uncover the molecular mechanism by which FAM111A suppresses vaccinia virus replication and demonstrate the mode of action of SPI-1 in antagonizing the antiviral function of FAM111A.

## Results

### FAM111A is a host restriction factor for VACV lacking SPI-1

To verify the antiviral effect of FAM111A, we cloned human FAM111A into a mammalian expression vector and expressed it ectopically in human A549 cells before infecting the cells with MVA, MVA engineered to express SPI-1 or/and C16 (MVA+SPI-1, MVA+SPI-1/C16), VACV-WR or VACV-ΔSPI-1. We examined the expression of viral proteins and viral replication by Western blotting analysis (0.3 PFU/cell) and virus quantification with plaque assay (0.03 PFU/cell), respectively. Transfected FAM111A was detectable with an anti-FAM111A antibody and overexpression of FAM111A dramatically decreased viral protein expression detected using an anti-VACV polyclonal antiserum (Fig. 1A). Notably, the decrease of viral protein expression was more phenomenal in VACV-ΔSPI-1 infected cells than in cells infected with VACV-WR or MVA+SPI-1/C16, suggesting SPI-1 may counteract with the function of FAM111A. In conformity with the results of Western blotting, over-expressing FAM111A significantly attenuated the replication of VACV in A549 cells and the effects were more pronounced for viruses that lack SPI-1 (VACV-ΔSPI-1 and MVA, Fig. 1B). The antiviral effect of FAM111A improved as we increased the amount of FAM111A transfected in MVA and VACV-ΔSPI-1-infected cells (Fig. 1C). To test the role of FAM111A on viral DNA replication, we examined the abundance of viral DNA using a set of primers for E11 in cells transfected with FAM111A and infected with VACV, MVA, and their recombinants indicated in the figure at 3 PFU/cell (Fig. 1D), and the blockage of viral DNA replication was observed in the presence of FAM111A^20^. In agreement with the above results, viral DNA loads were decreased significantly in both MVA and VACV-ΔSPI-1-infected cells in a FAM111A dose-dependent manner (Fig. 1E), further confirming the blockage of FAM111A on viral DNA replication.

**Figure 1.**
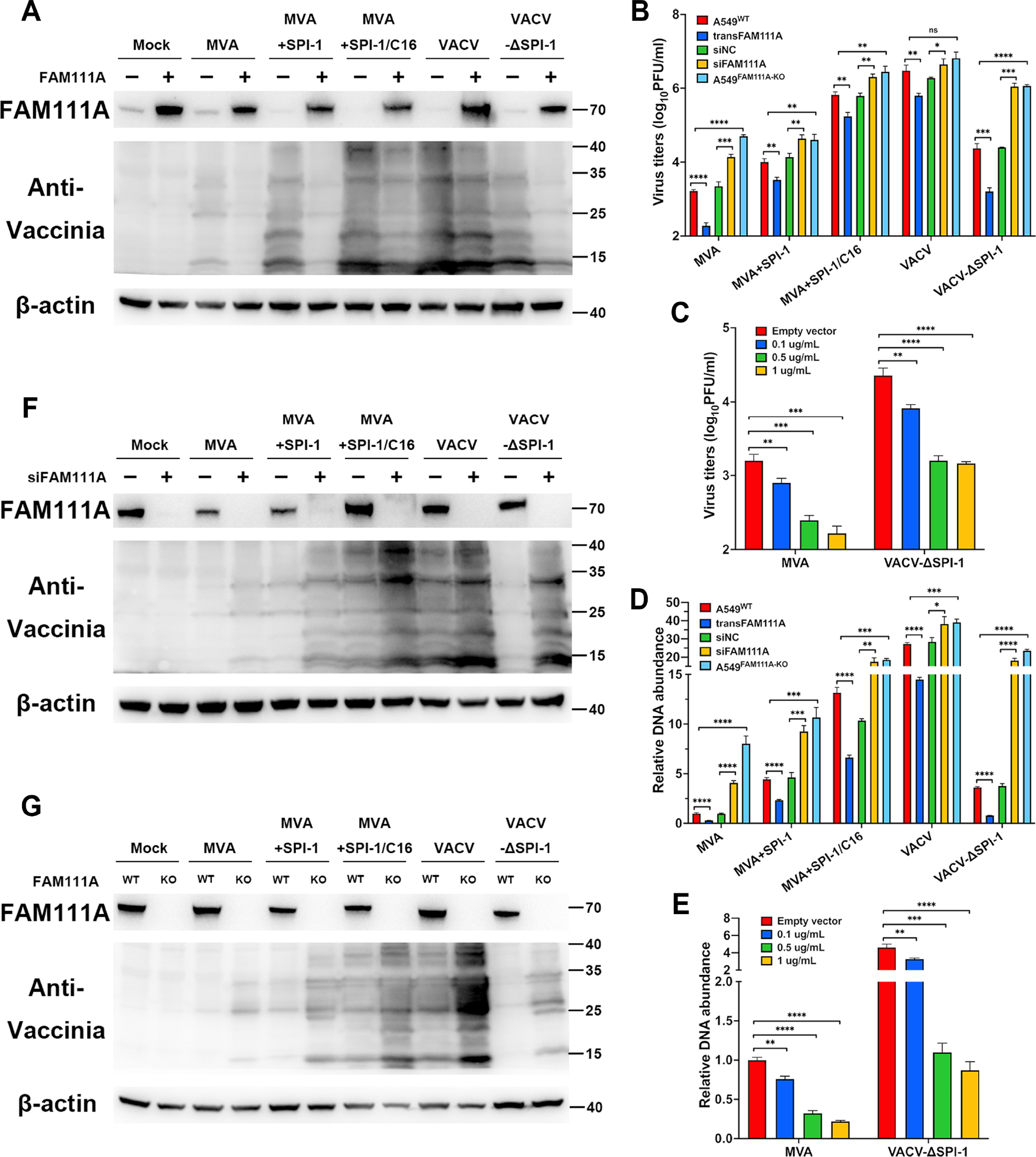
FAM111A prohibits the replication of vaccinia viruses lacking SPI-1 in A549 cells. **(A)** Human A549 cells were transfected with FAM111A or empty vector for 24 hours and infected with MVA, MVA+SPI-1, MVA+SPI-1/C16, VACV or VACV-ΔSPI-1 at 0.3 PFU/cell. Cell lysates were analyzed by SDS-PAGE and Western blotting with anti-FAM111A, anti- VACV, or anti-β-actin antibodies at 12 hpi. **(B and D)** Human A549 cells, A549 cells transiently transfected with FAM111A expression vector for 24 h, A549 cells transfected with siNC or siFAM111A for 48 hours, or A549^FAM111A-KO^ cells were infected with viruses described above at 0.03 PFU/cell. Viruses were harvested at 24 hpi and virus titers were measured by plaque assay on DF-1 (MVA, MVA+SPI-1, and MVA+SPI-1/C16) or BS-C-1 cells (VACV, VACV-ΔSPI-1). The bars represent the mean values of virus titers **(B)**; Indicated cells were infected with viruses described above at 3 PFU/cell. Total cellular DNA was harvested at 6 hpi and viral genomic DNA levels were determined by RT-qPCR using primers for E11 **(D)**. **(C and E)** Human A549 cells were transfected with a vector expressing FAM111A at concentrations of 0.1, 0.5, and 1 μg/mL for 24 hours and then infected with MVA or VACV-ΔSPI-1 at 0.03 PFU/cell. Viruses were harvested at 24 hpi and virus titers were determined by plaque assay using DF-1 cells (MVA) or BS-C-1 cells (VACV-ΔSPI-1). Bars represent the mean values of virus titers **(C)**; Indicated cells were infected with MVA or VACV-ΔSPI-1 at 3 PFU/cell. Total DNA was harvested at 6 hpi and viral genomic DNA levels were determined by RT-qPCR using primers for E11 **(E)**. **(F)** Human A549 cells were transfected with indicated siRNA for 48 hours and then infected with viruses described above at 0.3 PFU/cell. Cell lysates were analyzed by SDS-PAGE and Western blotting with anti-FAM111A, anti-VACV, or anti-β-actin antibodies at 12 hpi. **(G)** Human A549^WT^ or A549^FAM111A-KO^ cells were infected with viruses described above at 0.3 PFU/cell and cell lysates were collected and analyzed by SDS-PAGE followed by Western Blotting analysis with anti-FAM111A, anti-VACV or anti-β-actin antibodies at 12 hpi. Data in B-E represent the mean values ± SD (standard deviation) of 3 independent experiments. Data in A, F, and G are representative of 3 independent experiments. Statistics: ns, not significant; *, p < 0.05; **, p < 0.01; ***, p < 0.001; ****, p < 0.0001 by two-sided student’s t-test.

Next, we aimed to determine if depletion of endogenous FAM111A could relieve the replication of VACV lacking SPI-1. We first transiently knocked down the expression of FAM111A by transfecting A549 cells with siFAM111A or the negative control siNC. The expression of viral proteins determined by anti-VACV antiserum (Fig. 1F), virus titers (Fig. 1B), and viral DNA loads (Fig. 1D) were much higher in cells transfected with siFAM111A than in those transfected with only siNC as a negative control, and the comparison was more dramatic in MVA and VACV-ΔSPI-1-infected cells than in cells infected with SPI-1-expressing viruses. To further confirm our findings, A549^FAM111A-KO^ cell lines were generated using CRISPR-Cas9 technology with sgRNA targeting the 5’ region of FAM111A. The depletion of the protein was verified by Western blotting analysis (Fig. S1) and was further confirmed by Sanger sequencing. We infected A549^WT^ and A549^FAM111A-KO^ cells with the above-described viruses and examined the expression of viral protein expression, DNA abundance, and virus titers using the methods described. Depletion of FAM111A increased viral protein expression (Fig. 1G), virus titers (Fig. 1B), and viral DNA loads (Fig. 1D) dramatically. Replication of VACV-WT was however not affected by FAM111A depletion. Notably, even though VACV-ΔSPI-1 replicated better in FAM111A-KO cells than in WT cells, the titer was still lower than what VACV could achieve in either A549^WT^ or A549^FAM111A-KO^ cells, indicating additional restricting factors may exist for VACV-ΔSPI-1. To test if FAM111A is a species-specific host restriction factor, we transfected monkey BS-C-1 cells and avian DF-1 cells with human FAM111A and found human FAM111A failed to restrict virus replication in either of those cells, indicating the antiviral function of FAM111A may require other cellular factors that could be species-specific and are absent in non-human cells (Fig. S2). To inspect if viral infections influence FAM111A expression, proteins were collected from MVA- or VACV-infected cells for Western blotting analysis, and no change in FAM111A expression was observed post virus infection (Fig. S3) In summary, our results illustrated that FAM111A was a host restriction factor for VACV viruses that do not possess a functional SPI-1.

### Inhibition of VACV-ΔSPI-1 and MVA by FAM111A requires its functional protease domain

The C-terminal domain of FAM111A contains a trypsin-like motif that constitutes a catalytic triad consisting of His385, Aps439, and Ser541 and is highly conserved among mammals (Fig. S4)^21^. Mutation of S541A jeopardizes the enzymatic function of FAM111A and mutation of R569H is a hyperactive mutation found in patients with Kenny–Caffey syndrome (KCS) and osteocraniostenosis (OCS) exhibiting increased self-cleavage activity^21^. To assess if FAM111A’s antiviral effect is dependent on its protease activity, we generated vectors containing a loss-of-function mutant FAM111A^S541A,^ a hyperactive mutant FAM111A^R569H^ or a double mutant FAM111A^S541A+R569H^ that was previously shown to behave like a loss-of-function mutant (Fig. 2A). When transfected into human cells, FAM111A^S541A^ and FAM111A^S541A+R569H^ mutants caused accumulation of the protein comparing to FAM111A^WT^(Fig. 2B), a result of the decreased self-cleavage activity^7^. The hyperactive mutant FAM111A^R569H^ was expressed at a lower level compared to FAM111A^WT^, possibly due to excessive self-cleavage rather than proteasome-mediated degradation as the addition of MG132 (20 μM), a proteasome inhibitor, showed no effect on FAM111A protein levels (Fig. 2C). These results confirmed that the mutations indeed altered its protease activity. Next, A549 cells transfected with either FAM111A^WT^ or mutated FAM111As were infected with MVA or VACV-ΔSPI-1 and viral protein expression, virus growth, and DNA replication were monitored. The results showed that the reduction of VACV protein caused by overexpressing FAM111A was eliminated when the enzymatic activity was disrupted through S541 mutation, even though the mutants without a functional protease domain were expressed at much higher levels (Fig. 2D). It is worth noting that the hyperactive mutation FAM111A^R569H^ did not further decrease viral protein expression comparing to FAM111A^WT^, which may be the result of low protein accumulation due to excessive self-cleavage. In addition, FAM111As with loss-of-function mutations were no longer able to suppress virus replication (Fig. 2E) and viral DNA replication (Fig. 2F) in A549 cells, while the gain-of-function mutation did not show a significantly altered antiviral activity compared to FAM111A^WT^. As the PCNA interaction motif is required for its physiological function for host DNA replication, we generated a mutant that contained mutations in the PIP box (Fig. 2A) and found the loss of PCNA binding did not compromise the antiviral function of FAM111A (Fig 2G). These results demonstrated that the protease activity of FAM111A was indispensable for its antiviral function.

**Figure 2.**
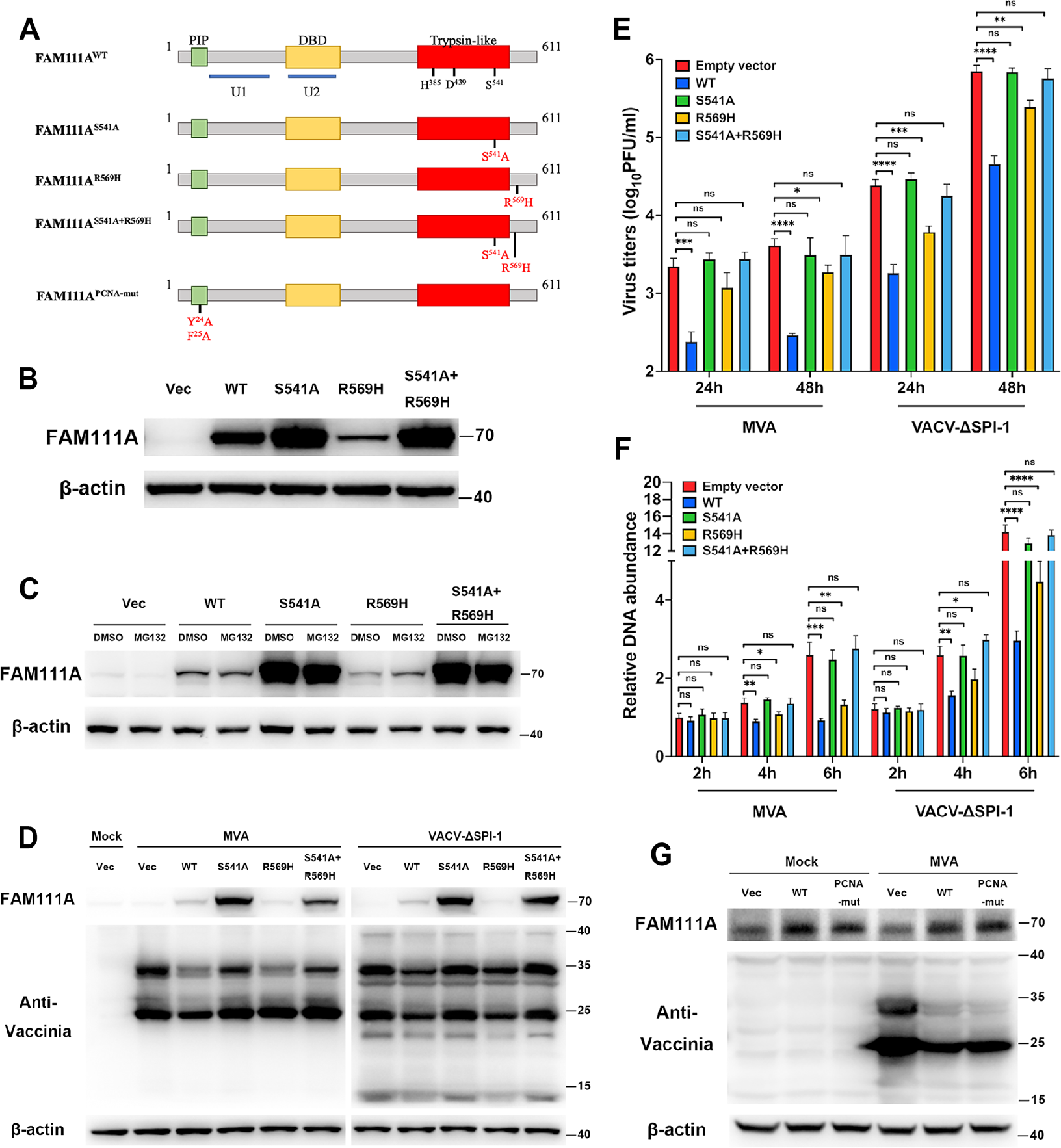
The antiviral effect of FAM111A is dependent on its protease activity. **(A)** Schematic of the protein structures of FAM111A with previously defined domains, the trypsin catalytic triplet mutants, and PCNA interaction domain mutant. Trypsin-like: the protease domain. PIP box: PCNA interaction domain. DBD: DNA binding domain. Catalytic residues in the protease domain are underlined. U1 and U2 are potential UBLs in FAM111A. **(B)** Human A549 cells transfected with the empty vector or vectors expressing FAM111A^WT^, FAM111^AS541A^, FAM111A^R569H,^ or FAM111A^S541A+R569H^ at the final concentration of 1 μg/mL. Cell lysates were collected and analyzed by SDS-PAGE followed by Western blotting analysis using anti-FAM111A or anti-β-actin antibodies. **(C)** Human A549 cells transfected with the empty vector or vectors expressing FAM111A^WT^, FAM111^AS541A^, FAM111A^R569H,^ or FAM111A^S541A+R569H^ at the final concentration of 1 μg/mL in the presence or absence of MG132 at 20 µM added 6h post-transfection. Cell lysates were collected and analyzed by SDS-PAGE followed by Western blotting analysis using anti-FAM111A or anti-β-actin antibodies. **(D-F)** Human A549 cells were transfected with an empty vector or vectors expressing FAM111A^WT^, FAM111^AS541A^, FAM111A^R569H,^ or FAM111A^S541A+R569H^ at 1µg/ml for 24 hours and then infected with MVA or VACV-ΔSPI-1 at 0.3 PFU/cell. Cell lysates were analyzed by SDS-PAGE and Western blotting with anti- FAM111A, anti-VACV, or anti-β-actin antibodies at 12 hpi **(D)**; Indicated cells were infected with MVA at 0.03 PFU/cell. Viruses were harvested at 24 hpi and virus titers were determined by plaque assay using DF-1 cells (MVA) or BS-C-1 cells (VACV-ΔSPI- 1). Bars represent the mean values of virus titers **(E)**; Indicated cells were infected with MVA or VACV-ΔSPI-1 at 3 PFU/cell. Total DNA was harvested at 6 hpi and viral genomic DNA levels were determined by RT-qPCR using primers for E11 **(F)**. **(G)** Human A549 cells were transfected with an empty vector or vectors expressing FAM111A^WT^ or FAM111A^PCNA-mut^ and then infected with MVA at 0.3 PFU/cell. Protein lysates were harvested and analyzed with SDS-PAGE and Western blotting analysis using anti- FAM111A, anti-VACV, and anti-β-actin antibodies. Data in E and F represent the mean values ± SD (standard deviation) of 3 independent experiments. Data in B-G are representative of 3 independent experiments. Statistics: ns, not significant; *, p < 0.05; **, p < 0.01; ***, p < 0.001; ****, p < 0.0001 by two-sided student’s t-test.

### VACV infection leads to the nuclear export of FAM111A through disruption of the nuclear pore complex (NPC)

Unlike most DNA viruses, poxvirus replication is restricted to the cytoplasm of infected cells. We thus deduce that FAM111A, a protein that existed in the nucleus under normal conditions, would have to be exported to the cytosol to exert its antiviral effect. To determine if viral infection triggers changes in the subcellular localization of FAM111A, A549 cells plated on coverslips were mock-infected or infected with MVA and the subcellular localization of FAM111A was determined by immunofluorescence using anti-FAM111A antibodies and Hoechst that could stain nucleus and virus factories. In mock-infected cells, FAM111A was exclusively found in the nucleus and the hyperactive mutation of FAM111A caused aberrant morphology of the nucleus, consistent with a previous report (Fig. 3A)^6^. When infected with MVA, however, FAM111A^WT^ was found primarily in the cytosol (Fig. 3B). In contrast, the two mutants with inactivated protease domain failed to translocate and were accumulated in the nucleus. FAM111A^R569H^, on the other hand, was also found predominantly in the cytosol but with a more concentrated distribution close to the viral factories (Fig. 3B). The subcellular localization of FAM111A and its mutants upon viral infection was also quantitated and the results were shown in Fig. 3C. We further isolated nuclear and cytosolic proteins and examined the abundance of FAM111A in each part and were able to confirm nuclear export of FAM111A upon MVA infection in a time-dependent manner (Fig. 3D-F). The above experiments were repeated in Hela cells and the results were consistent (Fig. S5).

**Figure 3.**
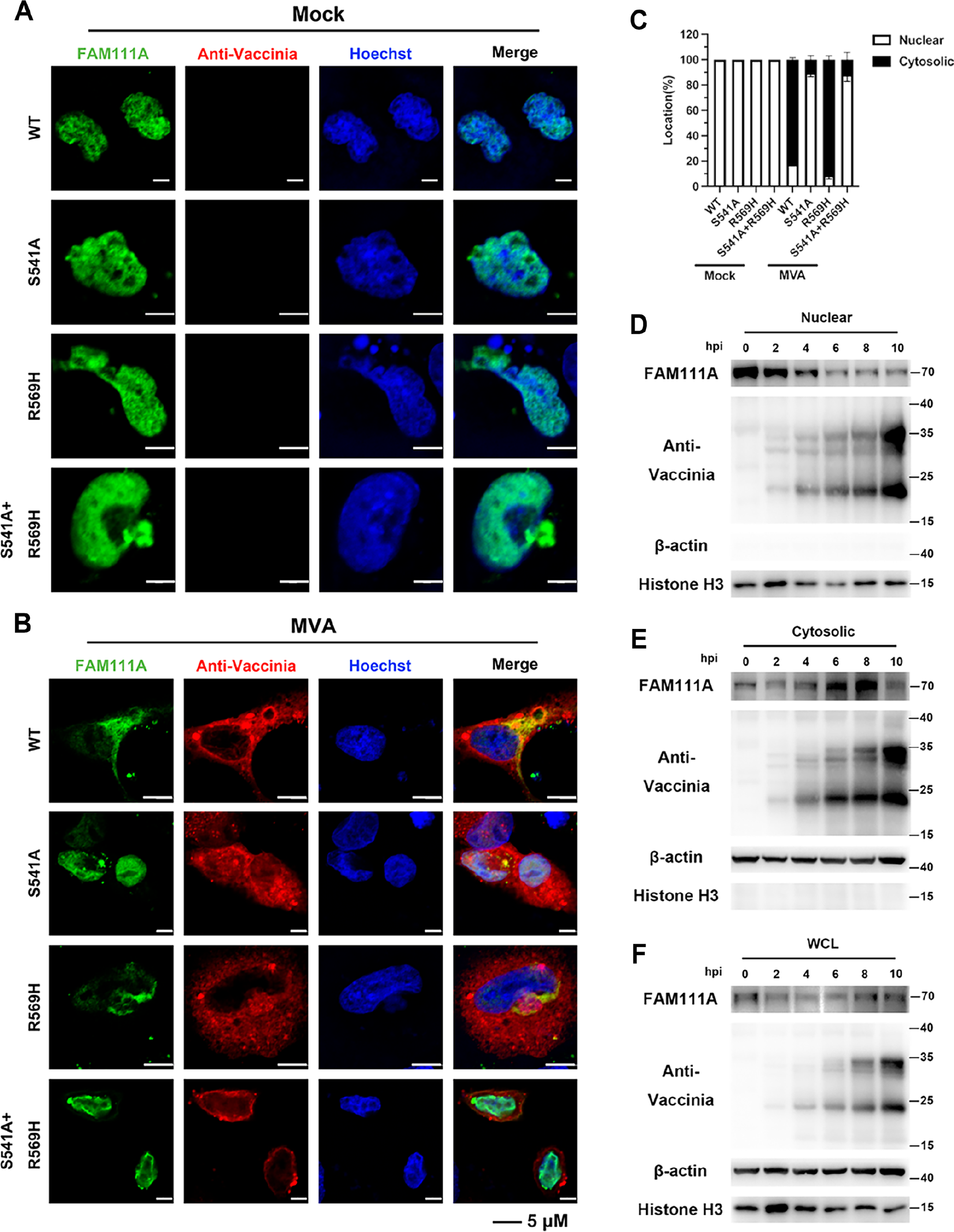
MVA infection prompts protease-dependent relocalization of FAM111A. **(A-C)** Human A549 cells plated on coverslips were transfected with vectors expressing FAM111A^WT^, FAM111^AS541A^, FAM111A^R569H,^ or FAM111A^S541A+R569H^ and then mock infected or infected with MVA at 3 PFU/cell. At 12 hpi, cells were then fixed, permeabilized, blocked, and stained with primary antibodies to FAM111A and VACV followed by fluorescent conjugated secondary antibodies. Hoechst was used to stain DNA **(A and B)**. The confocal analyses were performed in triplicates and the localization of FAM111A was quantified in 100 randomly selected cells and bars represent mean values +/- SD **(C)**. **(D-F)** Human A549 cells were infected with MVA at 3 PFU/cell and protein was harvested at time points indicated in the figure. Protein components in the nucleus and cytosol were separated by nuclear and cytoplasmic protein extraction kit and analyzed by SDS-PAGE and Western blotting analysis using anti-FAM111A, ati-VACV antibodies. Anti-β-actin and anti-histone H3 were used for cytosolic and nuclear loading controls, respectively. Data in C represent the mean values ± SD (standard deviation) of 3 independent experiments. Data in A, B, and D-F are representative of 3 independent experiments.

A previous report showed overexpression of FAM111A could destroy NPC by degrading NPC proteins^6^. We reasoned that nuclear export of FAM111A may be achieved through the destruction of the NPC by its protease activity. To test this hypothesis, A549 cells plated on coverslips were transfected with FAM111A or the loss-of-function and hyperactive mutants, and Mab414 was used to stain NPC for confocal microscopic analysis. As expected, transfection of FAM111A^WT^ and FAM111A^R569H^ disrupted normal NPC structure and FAM111A^S541A^ and FAM111A^S541A+R569H^ showed no such effect (Fig. S6A). In addition, NUP62, a core component of the NPC was found degraded by FAM111A (Fig. S6B). To test if endogenous FAM111A could degrade NPC upon viral infection, A549 cells were mock-infected or infected with MVA, MVA+SPI-1 at 3 PFU/cell for 6 hours and Mab414 antibody was used to examine NPC integrity by confocal microscopy. MVA infection led to disruption of the NPC in areas where FAM111A was concentrated (highlighted in the white box) and the nuclear membrane was undisturbed in mock-infected cells or cells infected with MVA+SPI-1 (Fig. 4A). The results from the confocal microscopic analysis were also quantitated as shown in Fig. 4B. These data revealed VACV infection triggered FAM111A-mediated NPC disruption and the subsequent nuclear export of FAM111A in the absence of SPI-1.

**Figure 4.**
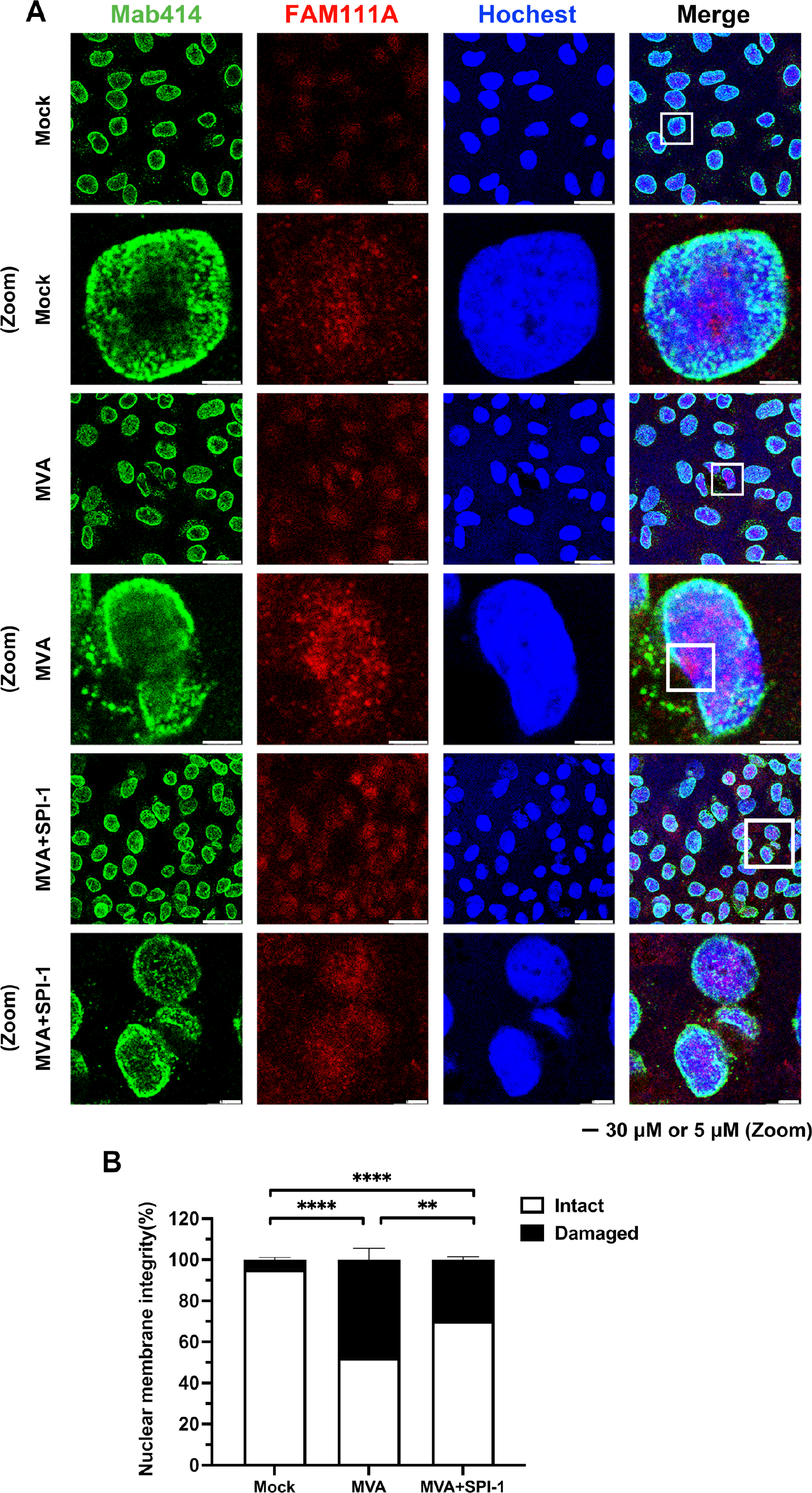
Virus infection induces NPC degradation by FAM111A. **(A-B)** Human A549 cells were mock infected or infected with MVA or MVA+SPI-1 at 3 PFU/cell. At 6 hpi, cells were then fixed, permeabilized, blocked, and stained with primary antibodies to Mab414 and FAM111A followed by fluorescent conjugated secondary antibodies. Hoechst was used to stain DNA **(A)**; The confocal analyses were performed in triplicates and the localization of FAM111A was quantified in 100 randomly selected cells and bars represent mean values +/- SD **(B)**. Data in B represent the mean values ± SD (standard deviation) of 3 independent experiments. Data in A and B are representative of 3 independent experiments.

### The cGAS-STING signaling pathway mediated the nuclear export of FAM111A upon VACV infection

As the lifecycle of poxvirus is complete in the cytosol of infected cells, we wondered how could FAM111A sense the invasion of VACV and initiate its migration to the cytosol. Cyclic GMP-AMP synthase (cGAS) is a crucial pattern recognition receptor activated by cytosolic DNA^22, 23^. To determine if cGAS is involved in FAM111A activation, we designed siRNA specific for cGAS and confirmed cGAS was knocked down efficiently (Fig. 5A). A549 cells transfected with siNC or sicGAS for 24 hours were left untreated or transfected with an empty vector, extracted genomic DNA from MVA, poly (dA:dT), a repetitive synthetic dsDNA, or poly (I:C), a molecule used to simulate dsRNA. Total protein was collected and components of nuclear and cytosolic proteins were isolated using the method described in figure 3. As shown in Fig 5A, sicGAS was able to repress the expression of cGAS and phosphorylation of STING but had no effect on the expression of FAM111A, total STING, PCNA, or histone H3. Transfection of plasmid DNA, viral genomic DNA, or poly (dA:dT) all led to increased nuclear export of FAM111A while the transfection of poly (I:C) did not (Fig. 5C). Poly (dA:dT) showed strong induction of FAM111A expression and the reason was unknown. Importantly, the cytosolic accumulation of FAM111A caused by transfection of plasmid DNA or viral genomic DNA was largely reduced in cells transfected with sicGAS, indicating nuclear export of FAM111A was induced by dsDNA in the cytosol through the cGAS-STING signaling pathway.

**Figure 5.**
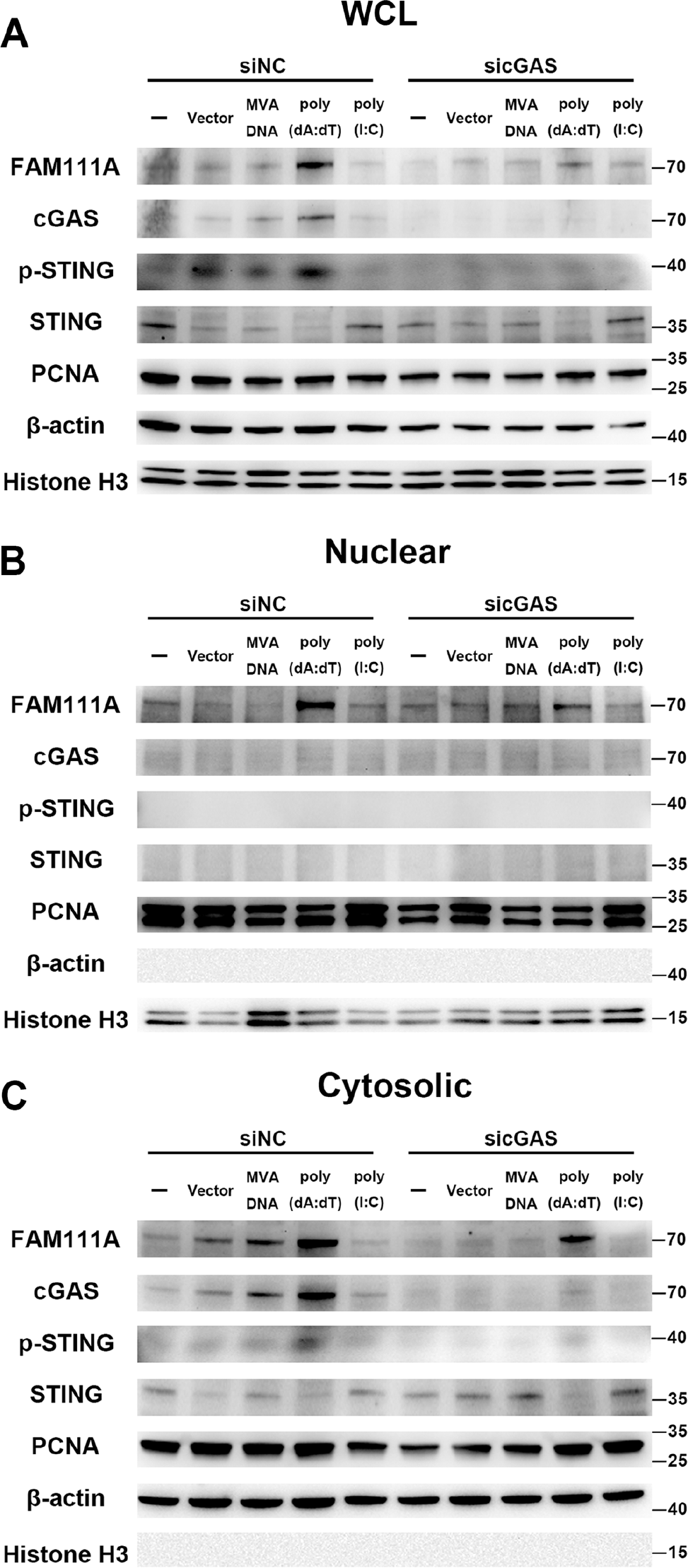
The relocalization of FAM111A is enabled by the activation of the cGAS-STING pathway. Human A549 cells were transfected with siNC or sicGAS for 48 hours and then transfected with empty vector, DNA collected from MVA virus particles, poly (dA:dT), or poly (I:C) for 8 hours. Nuclear and cytosolic proteins were separated using nuclear and cytoplasmic protein extraction kits and analyzed by SDS-PAGE and Western blotting analysis using anti-FAM111A, anti-cGAS, anti-p-STING, anti-STING, anti-PCNA, anti-β- actin, and anti-histone H3 antibodies. Anti-β-actin and anti-histone H3 were used for cytosolic and nuclear loading controls, respectively. Data are representative results of 3 independent experiments.

### FAM111A targets virus DNA binding protein I3 for degradation

The above-mentioned experiments demonstrated that VACV infection induced nuclear export of FAM111A through the cGAS signaling pathway and the export of FAM111A was achieved through NPC destruction. It was hence important to investigate how FAM111A exerted its antiviral activity. We first hypothesized that FAM111A could directly bind to and catalyze the degradation of certain viral proteins via its active protease domain. We designed an experiment to pull down the viral interactome of FAM111A in virus-infected cells and identified its components with mass spectrometric analysis (Fig. 6A). In this assay, VACV I3 showed the most significant difference in protein abundance between FAM111A-transfected and vector-transfected cells and was further investigated as a putative target of FAM111A (Fig. 6B). Vaccinia virus I3 contains a single-stranded DNA binding domain and is essential for virus DNA replication, deletion of I3 did not affect viral early protein synthesis or core disassembly but blocked DNA replication ^24, 25^. The interaction between FAM111A and I3 was first confirmed by co-immunoprecipitation (co-IP) analysis by using either FAM111A or I3 as the bait (Fig. 6C-D). To determine if the interaction was DNA-dependent as FAM111A contains a double-stranded DNA binding domain (DBD), cell lysates were incubated with or without 250U/ml of benzonase, an endonuclease known to degrade all forms of DNA without base preference. The association of I3 and FAM111A disappeared when benzonase was added (Fig. 6C-D). As FAM111A contains a DBD and no discernible RNA binding motifs, we concluded that the interaction between FAM111A and I3 was DNA-dependent. Additional experiments were carried out to validate the co-localization of I3 and FAM111A within cells (Fig. 6E). A549 cells on coverslips were co-transfected with vectors expressing viral I3 and human FAM111A for confocal microscopic analysis. As expected, I3 was associated with Hoechst-stained virus factories in the cytoplasm of infected cells and the majority of FAM111A colocalized with I3 (Fig. 6E). In summary, the association of FAM111A and I3 was shown by immunoaffinity purification and further suggested by their intracellular colocalization in the cytoplasm.

**Figure 6.**
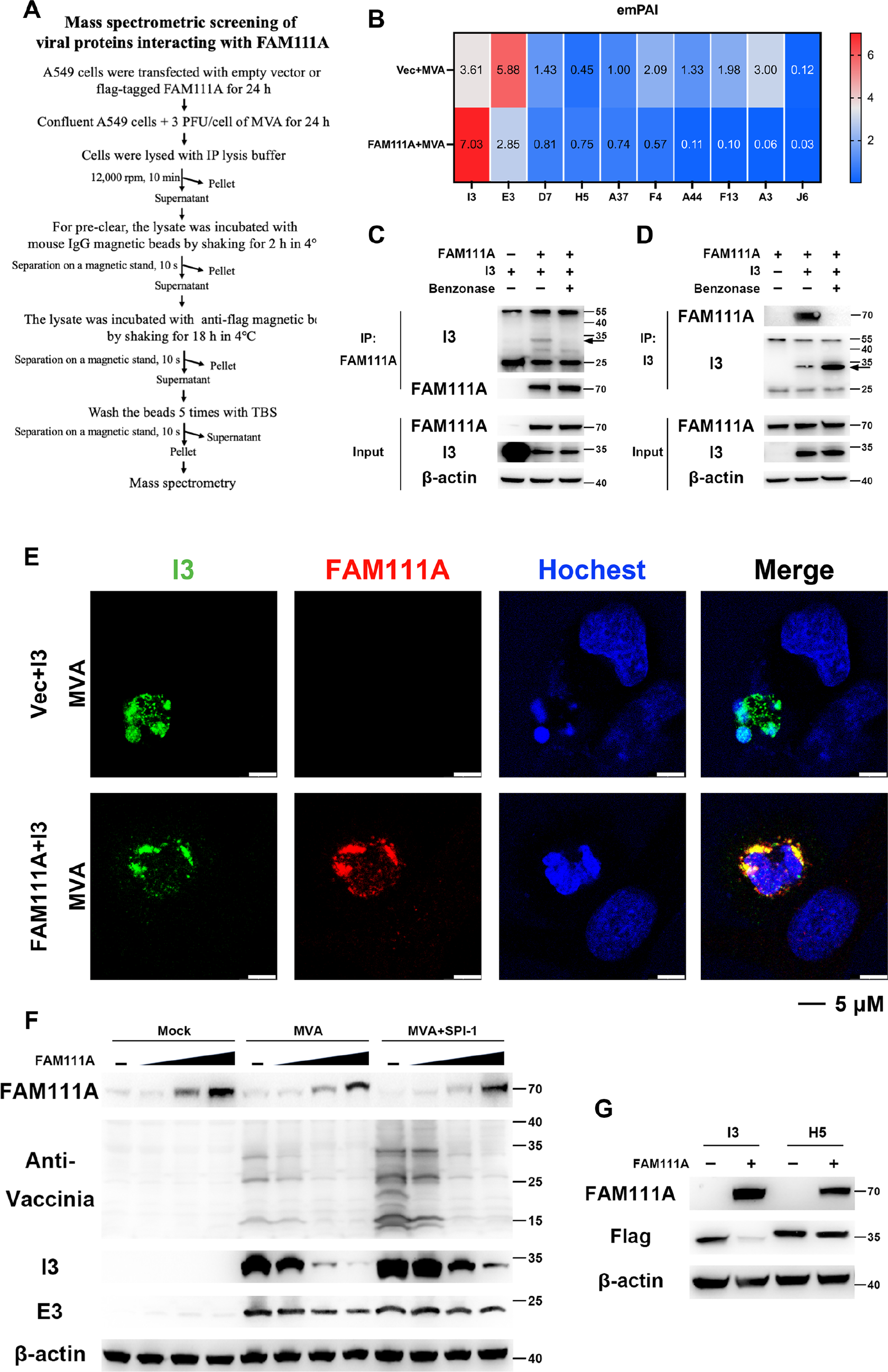
FAM111A mediates the degradation of viral I3 in a DNA-dependent manner **(A)** Schematic flow chart of mass spectrometric screening of viral proteins interacting with FAM111A. **(B)** Mass spectrometry was used to identify the MVA viral proteins that interacted with FAM111A, and the relative content of different viral proteins was expressed by the exponentially modified protein abundance indice (emPAI). **(C and D)** Human A549 cells were transfected with vectors expressing myc-tagged FAM111A or flag-tagged I3 for 24 hours. Cell lysates were first incubated with or without Benzonase (250 U/mL) at 37°C for 20 min and then incubated with control magnetic beads or myc-/flag- conjugated beads at 4°C for 18 hours. Beads were extensively washed and proteins were eluted with SDS-loading buffer and analyzed by SDS-PAGE and Western blotting analysis. The arrow indicates the correct band size for I3, and the light chain or heavy chain antibody bands are at 25 kDa or 55 kDa, respectively. **(E)** A549 cells plated on coverslips were co-transfected with VACV-I3 and FAM111A for 24 hours and then infected with MVA at 3 PFU/cell. At 12 hpi, cells were then fixed, permeabilized, blocked, and stained with primary antibodies to I3 and FAM111A followed by fluorescent conjugated secondary antibodies. Hoechst was used to stain DNA. Scale bars were shown at the bottom-right corners. **(F)** Human A549 cells were transfected with a vector expressing FAM111A at concentrations of 0.1, 0.5, and 1 μg/mL for 24 hours and then infected with MVA or MVA+SPI-1 at 0.3 PFU/cell. Cell lysates were analyzed by SDS- PAGE and Western blotting with anti-FAM111A, anti-VACV, anti-I3, anti-E3, or anti-β-actin antibodies at 24 hpi. **(G)** Human A549 cells were co-transfected with flag-tagged I3 or H5 and vectors expressing FAM111A for 24 hours. Cell lysates were analyzed by SDS-PAGE and Western blotting analysis using anti-FAM111A, anti-flag, and anti-β-actin antibodies. Data in C-G are representative of 3 independent experiments.

Next, experiments were performed to investigate if FAM111A resulted in virus I3 degradation. A549 cells transfected with increasing amounts of FAM111A were mock-infected or infected with MVA or MVA+SPI-1 at 3 PFU/cell and virus protein expression was examined by Western blotting analysis. Levels of I3 reduced as the amounts of FAM111A transfected increased in infected cells (Fig. 6F). It is worth noting that FAM111A was also able to lessen I3 abundance in MVA+SPI-1-infected cells. One possible explanation is that the amount of SPI-1 present was not sufficient to fully restrict FAM111A when it was transfected at a high concentration. Nevertheless, it was apparent that I3 in MVA+SPI-1-infected cells appeared more resistant to FAM111A than that in MVA-infected cells (Fig. 6F). Importantly, E3, a viral early protein that binds dsRNA, was insensitive to the presence of FAM111A (Fig. 6F). To further verify the specificity of I3 degradation by FAM111A, we co-transfected A549 cells with FAM111A and flag-tagged I3 or H5, another DNA binding protein, and found like E3, H5 was unaffected by FAM111A (Fig. 6G). These results demonstrated FAM111A was specifically associated with virus DNA binding protein I3 and mediated its degradation upon virus infection.

### FAM111A promotes I3 degradation through autophagy but not its peptidase activity

To investigate if FAM111A directly degrades virus I3 through its C-terminal protease activity, we simultaneously co-transfected A549 cells with VACV I3 and FAM111A or its hyper- and hypo-active mutants and monitored protein levels of I3. Surprisingly, I3 was degraded when not only FAM111A^WT^ but also when FAM111A mutants with inactivated (FAM111A^S541A^ and FAM111A^S541A+R569H^) protease activity were transfected (Fig. 7A). As we previously discovered that the protease activity of FAM111A was crucial for its nuclear export, these combined results suggested that the FAM111A mutants with inactivated peptidase domain may bypass nuclear export and associate with I3 directly when they were co-transfected. The co-transfection of FAM111A and I3 may lead to a pre-mature association of FAM111A and I3 in the cytoplasm and result in I3 degradation. To determine whether the subcellular localization of FAM111A could affect its antiviral function, we decided to alter our strategy by transfecting cells with FAM111A or its derivative mutants for 24 hours before transfecting I3. By doing this, FAM111A expressed could be localized to the correct subcellular position before interacting with I3 (Fig. 7B). In addition, we manipulated the subcellular localization of FAM111A by either attaching a nucleus export signal (NES) or disrupting its nucleus localization signal (NLS-mut) (Fig. 7C). A549 cells were transfected with FAM111A^WT^, FAM111A^DBD-mutant^, FAM111A^S541A^, FAM111A^S541A+NES,^ or FAM111A^NLS-mut^ for 24 hours, and then transfected with I3 and the levels of FAM111A and I3 were examined by Western blotting analysis. As expected, mutants with disrupted double-stranded DNA binding domain or inactivated protease domain abolished their ability to degrade I3 (Fig. 7D) and were found primarily in the nucleus (Fig. S7). Remarkably and interestingly, the mutant with an inactivated protease domain combined with an NES motif, which was found mostly in the cytosol, colocalized with I3 (Fig. S7), and led to its degradation (Fig. 7D). Similarly, the mutant with a disrupted NLS was found primarily in the cytoplasm (Fig. S7) and was also able to mediate I3 degradation. These results demonstrated that the protease domain of FAM111A was only required for nuclear export but not I3 degradation.

**Figure 7.**
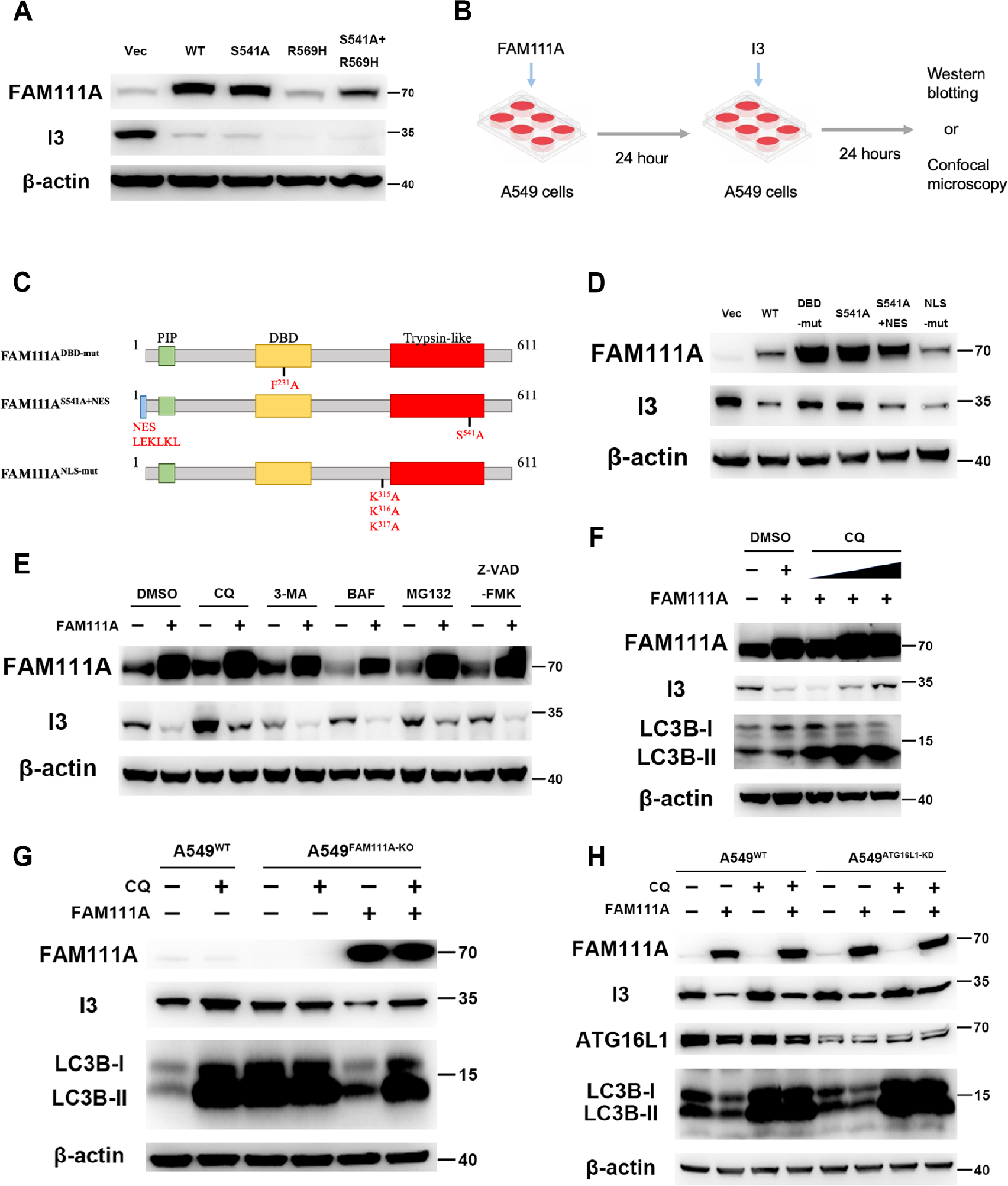
Cytosolic FAM111A promotes I3 degradation through autophagy. **(A)** Human A549 cells were co-transfected with VACV-I3 and vectors expressing FAM111A^WT^ and lab-generated mutants indicated in figure 2(A) for 24 hours. Cell lysates were analyzed by SDS-PAGE and Western blotting analysis using anti-FAM111A, anti-I3, and anti-β-actin antibodies. **(B)** Schematic flowchart of transfection experiments. Human A549 cells were transfected with vectors expressing FAM111A for 24 hours before I3 transfection. After 24 hours, cells were analyzed by Western blotting or confocal microscopy. **(C)** Diagram of structures of FAM111A DNA binding domain mutant, FAM111A^S541A^ mutant with nuclear export signal (NES), and FAM111A nuclear localization signal mutant (NLS-mut). **(D)** Human A549 cells were transfected with vectors expressing FAM111A^WT^ and lab-generated mutants indicated in the figure for 24 hours and then transfected with I3. After 24 hours, cell lysates were analyzed by SDS- PAGE and Western blotting analysis using anti-FAM111A, anti-I3, anti-LC3B, and anti-β- actin antibodies. **(E)** Human A549 cells were transfected with an empty vector or vector expressing FAM111A for 24 hours before transfecting with VACV-I3. Different drugs were added at the time of I3 transfection. After 24 hours, cell lysates were analyzed by SDS- PAGE and Western blotting analysis using anti-FAM111A, anti-I3, and anti-β-actin antibodies. **(F)** Human A549 cells were transfected with an empty vector or vector expressing FAM111A for 24 hours before I3 transfection. CQ at various concentrations was added. After 24 hours, cell lysates were analyzed by SDS-PAGE and Western blotting analysis using anti-FAM111A, anti-I3, anti-LC3B, and anti-β-actin antibodies. **(G)** Human A549^WT^ or A549^FAM111A-KO^ cells were transfected with an empty vector or vector expressing FAM111A for 24 hours before I3 was transfected and CQ added. After 24 hours, cell lysates were analyzed by SDS-PAGE and Western blotting analysis using anti-FAM111A, anti-I3, anti-LC3B, and anti-β-actin antibodies. **(H)** Human A549^WT^ or A549^ATG16L1-KD^ cells were transfected with an empty vector or vector expressing FAM111A for 24 hours before I3 transfected. DMSO or CQ was added at the time of I3 transfection. After 24 hours, cell lysates were analyzed by SDS-PAGE and Western blotting analysis using anti-FAM111A, anti-I3, anti-ATG16L1, anti-LC3B, and anti-β-actin antibodies. Data in A and D-H are representative of 3 independent experiments.

Next, we explored the nature of I3 degradation by using a set of chemical inhibitors targeting various cellular pathways known to modulate protein degradation. A549 cells were sequentially transfected with FAM111A and I3 (Fig. 7B) in the presence or absence of chemicals inhibiting autophagy [10 μM chloroquine (CQ), 5 mM 3-methyladenine (3-MA), 0.5 μM bafilomycin A1 (BAF)], proteasomal degradation (20 μM MG132) or pan-caspase activity (20 μM Z-VAD-FMK). While the addition of CQ and MG132 both alleviated FAM111A-induced degradation of I3, the addition of 3-MA, BAF or Z-VAD-FMK showed no discernible effects (Fig. 7E). Proteasome-mediated I3 degradation was ruled out as I3 co-purified with FAM111A exhibited no sign of ubiquitination (Fig. S8). The result of MG132 was likely due to an unknown off-target effect. We next performed a dose-dependent experiment and further confirmed that the addition of CQ caused the accumulation of LC3B-II and led to the rescue of FAM111A-mediated I3 degradation (Fig. 7F). Moreover, CQ slightly increased the expression I3 when cells were transfected with only I3 but not FAM111A (Fig. 7G), this was possibly due to I3 degradation stimulated by endogenous FAM111A, as this phenomenon was not observed in A549^FAM111A-KO^ cells (Fig. 7G). To rule out the possibility that blockage of I3 degradation was due to the off-target effect of CQ, we manipulated the level of ATG16L1, a key regulator of autophagic flow, by generating an A549 cells line in which the endogenous level of ATG16L1 was stably knocked down by shRNA and named it A549^ATG16L1-KD^ (Fig. 7H). Similar to the effect observed with CQ, depression of ATG16L1 was able to rescue FAM111A-mediated I3 degradation, suggesting the role of autophagy in such a process. The above results showed that FAM111A interacted with virus I3 and promoted I3 degradation by utilizing the autophagy pathway, rather than its protease activity.

### Poxvirus SPI-1 blocks the nuclear export of FAM111A through its serpin activity

SPI-1 (C12 in VACV-cop) has previously been shown to function as a host range factor for MVA^18^. To explore its role in FAM111A inhibition, we first observed its subcellular localization by infecting FAM111A-transfected A549 cells with MVA or MVA+SPI-1 at 3 PFU/cell and probed SPI-1 using a myc antibody at 12 hpi as the SPI-1 inserted into MVA contains a myc tag at its N-terminus^18^. The confocal microscopic analysis showed that SPI-1 was mainly localized in the nucleus during MVA+SPI-1 infection (Fig. S9). In addition, nuclear export of FAM111A activated by MVA was no longer observed when cells were infected with MVA+SPI-1 (Fig. 8A), suggesting SPI-1 prevented FAM111A from exporting to the cytoplasm. To investigate if SPI-1 was associated with FAM111A, we performed immunoaffinity purification using either SPI-1 or FAM111A as the bait and observed a strong interaction between FAM111A and SPI-1 (Fig. 8B-C). As FAM111A restricted viral replication by degrading virus I3, we next inspected if SPI-1 could interfere with FAM111A-mediated I3 degradation. A549 cells were co-transfected with FAM111A, VACV I3, and VACV-SPI-1. Consistent with previous results, transfection of FAM111A led to a prominent I3 degradation when SPI-1 was absent, however, this effect was alleviated when plasmid expressing VACV-SPI-1 was co-transfected (Fig. 8D). Interestingly, SPI-1 from MPXV, a close relative of VACV, and lumpy skin disease virus, a member of the capripoxviruses, displaying 96.4% and 37.8% sequence identity with VACV SPI-1, respectively, could also inhibit FAM111A-mediated I3 degradation. A previous report identified cathepsin G as one of the cellular targets of rabbitpox virus SPI-1 and showed mutations in the reactive-site loop (RSL) of SPI-1 abolished its interaction with cathepsin G^19^. We thus introduced the two mutations T305R, F318A separately or in combination into VACV SPI-1 and examined their effects on FAM111A inhibition (Fig. 8E). We used the same transfection strategy described in Fig. 7B by transfecting FAM111A 24 hours before transfecting I3 and SPI-1. As shown in Fig. 8E, wt-SPI-1 was able to rescue FAM111A-mediated I3 degradation, however, the SPI-1 mutants with mutations in the RSL region failed to do so even though expressed at comparable levels with the wt-SPI-1. These results suggested that SPI-1 interacted with FAM111A via the RSL region and demonstrated that SPI-1 antagonized FAM111A’s antiviral function by inhibiting the serine protease activity of FAM111A.

**Figure 8.**
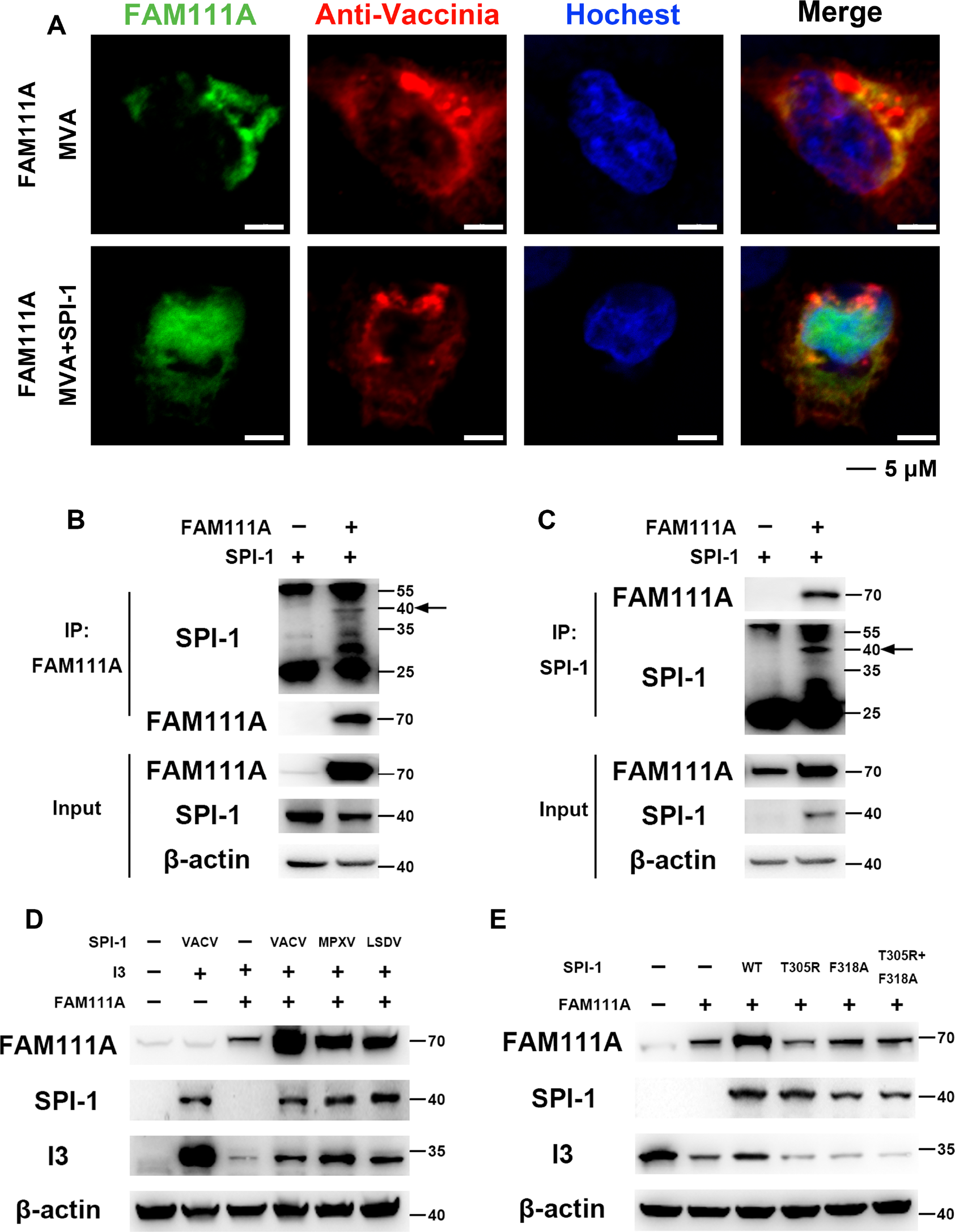
Poxvirus SPI-1 antagonizes nuclear export and I3-degradation of FAM111A. **(A)** Human A549 cells were transfected with an empty vector or vector expressing FAM111A for 24 hours and then infected with MVA or MVA+SPI-1 at 3 PFU/cell. After 12 hours, the cells were then fixed, permeabilized, blocked, and stained with primary antibodies to FAM111A and VACV followed by fluorescent conjugated secondary antibodies. Hoechst was used to stain DNA. Scale bar at bottom. **(B and C)** Human A549 cells were transfected with vectors expressing myc-tagged FAM111A or flag-tagged SPI- 1 for 24 hours. Cell lysates were incubated with control magnetic beads or myc-/flag- conjugated beads at 4°C for 18 hours. Beads were extensively washed and proteins were eluted with SDS-loading buffer and analyzed by SDS-PAGE and Western blotting analysis. The arrow indicates the correct band size for SPI-1, and the light chain or heavy chain antibody bands are at 25 kDa or 55 kDa, respectively. **(D)** Human A549 cells were co-transfected with flag-tagged SPI-1 of different viruses and a vector expressing FAM111A for 24 hours before I3 transfection. After 24 hours, cell lysates were analyzed by SDS-PAGE and Western blotting analysis using anti-FAM111A, anti-flag, anti-I3, and anti-β-actin antibodies. **(E)** Human A549 cells were co-transfected with flag-tagged SPI-1 of different mutants and a vector expressing FAM111A for 24 hours. Then indicated cells were infected with MVA at 0.3 PFU/cell. After 24 hours, cell lysates were analyzed by SDS-PAGE and Western blotting analysis using anti-FAM111A, anti-flag, anti-I3, and anti- β-actin antibodies. Data in A-E are representative of 3 independent experiments.

We summarized our findings in a diagram presented in Fig. 9.

**Figure 9.**
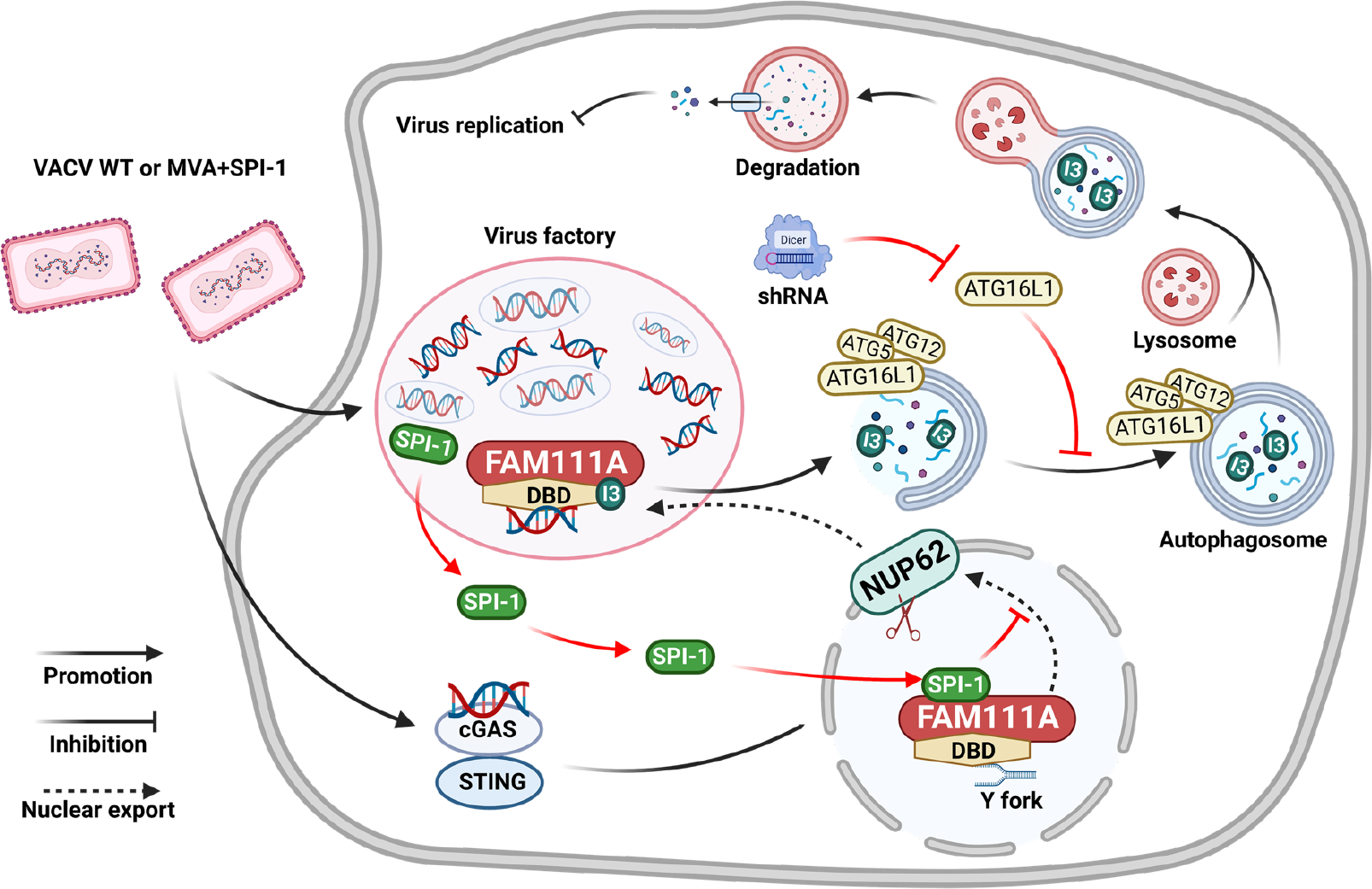
FAM111A inhibits viral replication by inducing I3 degradation and is antagonized by VACV SPI-1. After SPI-1-deficient MVA or VACV invaded cells, the viral genomic DNA was recognized by the cGAS-STING signaling pathway, and its downstream signal activated nuclear export of FAM111A, allowing FAM111A in the nucleus to enter the cytoplasm by degrading the NPC. FAM111A is then recruited to the virus factory and binds to viral protein I3, which is thus degraded by autophagy. VACV SPI-1 enters the nucleus and inhibits the serine protease activity of FAM111A to prohibit its antiviral effect.

## Discussion

FAM111A has been postulated to be a serine protease regulated in a cell-cycle-dependent fashion and conserved in mammals^7^. Recent studies showed FAM111A may restrict SV40 HR mutants and vaccinia virus infections^6, 8^. We examined the effect of overexpressing or depression of FAM111A on virus replication using VACV that lacks a functional SPI-1 (VACV-ΔSPI-1) or MVA, which lost its SPI-1 during repeated passages in avian cells. Deletion of FAM111A with CRISPR-Cas9 or transient suppression with RNAi enhanced VACV-ΔSPI-1 or MVA replication in human A549 cells. In addition, we discovered that its antivral effect occurred post-replicatively as a viral early gene E3 was expressed in the presence of FAM111A. Moreover, transient transfection of FAM111A in human cells exhibited a strong inhibitory effect on viral DNA replication, viral protein expression, and virus replication, further confirming the antiviral function of FAM111A. The trypsin-like domain of FAM111A contains a catalytic triad including His385, Aps439, and Ser541 and a DNA-binding domain that plays essential roles for its localization at the replication forks^5, 7, 26, 27^. FAM111A was also shown to bind to the large T antigen of SV40 and inhibit the formation of the replication center of SV40 mutants^8^. However, the replication of VACV differs from that of SV40 as the former completes its entire replication cycle in the cytoplasm of infected cells. As a result, nuclear export is the first step for FAM111A to confer its antiviral effect for VACV. Mutations of the catalytic triad of FAM111A abolished its antiviral activity for VACV-ΔSPI-1 and MVA as disruption of NPC no longer occurred (Fig. S6). In addition, transfection of FAM111A with an unmutated or hyperactive peptidase domain disrupted the nuclear membrane, which was previously reported and further demonstrated that the exportation of FAM111A upon infection is dependent on its peptidase domain^6^. Following MVA infection, FAM111A translocated from the nucleus to the cytoplasm and co-localized with viral factories stained by Hoechst or I3 (Fig. 3 and 6). The change in FAM111A’s subcellular localization was dependent on its DNA-binding domain but not the PCNA-interaction motif. Another interesting question was how FAM111A, a protein exclusively found in the nucleus, senses VACV infection in the cytosol. Panda et al. postulated that replication factor C (RFC) 1-5 could load PCNA and may act as a DNA sensor recruiting FAM111A to viral genomes^9^. Nonetheless, we found that mutations in the PCNA-interecting motif (FAM111A^PCNA-mut^) did not influence VACV restriction, suggesting the recruitment of FAM111A to the viral factory is not PCNA mediated. Rather, mutations in the DBD abolished the nuclear export of FAM111A and resulted in its nuclear accumulation, indicating the involvement of DNA binding in FAM111A recruitment (Fig.S7). This was further validated by the finding that FAM111A^DBD-mut^ no longer promoted I3 degradation. Interestingly, the reduction of cGAS expression in cells inhibited the nuclear export of FAM111A, signifying the existence of a regulatory axis that associated cGAS-STING signaling pathway with FAM111A (Fig. 5). More needs to be done to clarify components in the cGAS-FAM111A regulatory pathway and to explore the nature of FAM111A activation upon virus infection. Previous studies showed that infection of an SV40 host range mutant enhanced the protease activity of FAM111A, leading to impairment of nuclear barrier function, cell morphology, and normal distribution of the nuclear pore^6^. These results were in line with our observation but the upstream activating signal for FAM111A may be different as SV40 replicates in the nucleus.

Using an immunoprecipitation assay coupled with mass-spectrometry, we were able to identify the viral binding partners of FAM111A during MVA infection. We discovered that cytosolic FAM111A interacted with viral I3 in a DNA-dependent manner. VACV I3 is a 34 kDa phosphoprotein essential for poxvirus genome replication, VACV-ΔI3 showed normal early gene expression and core disassembly but reduced DNA replication in non-complementing cells, a phenotype resembles that of FAM111A overexpression^25, 28^. Like FAM111A, I3 can also bind DNA with its single-stranded DNA binding domain (SSB)^29^. The interaction between FAM111A and I3 was terminated with benzonase treatment, and the FAM111A mutant that lacks a functional DNA binding domain no longer displayed any antiviral activity (Fig. 6 and 7). These observations highlight the importance of DNA binding in its interaction with I3 and the recruitment of FAM111A to virus factories. Cytoplasmic FAM111A led to efficient I3 degradation, obstructing virus DNA replication and subsequent virus replication. Surprisingly, disease-associated mutations or loss-of-function mutations of FAM111A did not influence its ability to promote I3 degradation as long as they were expressed in the cytoplasm. By manipulation of the subcellular localization of FAM111A using NES or interruption of NLS signals within FAM111A, we found that the catalytic activity of FAM111A was only required for nuclear export but not I3 degradation as the FAM111A^S541A^ containing an NES (FAM111A^S541A+NES^) was still able to induce I3 degradation (Fig. 7). Moreover, we found the degradation of I3 by FAM111A did not rely on proteasomes, nor did it use the caspases. Instead, CQ treatment or depression of ATG16L1 by shRNA were both able to relax the degradation of I3 by FAM111A, demonstrating the role of autophagy in I3 degradation. In our assay, 3-MA and bafilomycin A1, two other inhibitors of autophagy failed to negate FAM111A-mediated I3 degradation. The exact reason was unknown but we speculated that it may be because bafilomycin A1, 3-MA, and chloroquine inhibit different steps of the autophagic flow and VACV may possess strategies to compensate for the blockade of autophagosome nucleation and lysosomal acidification, which were inhibited by 3-MA and bafilomycin A1, respectively^30^.

VACV SPI-1 encoded by the C12 gene in RPXV was previously thought to affect the virus host range by inhibiting the serine protease activity of cathepsin G^19^. Vaccinia virus lacking SPI-1 exhibited low late and intermediate mRNAs but undiminished early mRNA^31^.

Deletion of SPI-1 did not affect virus virulence in a murine intranasal model^32^. A more recent study confirmed the role of SPI-1 as a host range factor for the replication of MVA in human cells^18^. We found SPI-1 mainly distributed in the nucleus after virus infection and interacted with FAM111A in a co-IP assay. The presence of SPI-1 was able to prevent FAM111A’s nuclear export, thereby antagonizing its antiviral function. In RPXV, SPI-1 was predicted to be associated with viral proteins essential for DNA replication^33^. Our results were in agreement with this hypothesis as we observed interactions and co-localization of SPI-1 and FAM111A within the nucleus of infected cells. Furthermore, we found that SPI-1 proteins from different poxviruses [VACV, monkeypox virus (MPXV), and lumpy skin disease virus (LSDV)] all inhibited FAM111A-mediated I3 degradation. Importantly, mutations of the two residues in the RSL of SPI-1 (T305R, F318A) abrogated its function in mitigating FAM111A-promoted I3 degradation without significantly affecting SPI-1 expression. As these two residues were previously reported to be essential for cathepsin G binding, we speculated that SPI-1 interacted with FAM111A via a similar region in its RSL^19^.

As far as we know, our work is the first report of a mechanism by which FAM111A prohibited cytoplasmic DNA virus replication (summarized in Fig. 9). Our results indicated that nuclear components normally involved in mammalian cell DNA replication could translocate from the nucleus to the cytoplasm and restrict VACV replication by degrading key viral DNA replication-regulating protein I3 using the autophagic machinery. In addition, we presented an explanation for the immune evasion function of VACV SPI-1.

## Materials and methods

**Cell Culture and Generation of Stable Cell Lines.** Human A549 cells (ATCC CCL-185) were purchased from ATCC and maintained in Dulbecco’s Modified Eagle Medium (DMEM)/F-12 (Gibco) supplemented with 10% fetal bovine serum (FBS, Gibco) and 1% penicillin-streptomycin solution (Solarbio). HeLa cells (ATCC CCL-2), 293T cells (ATCC CRL-3216), and DF-1 cells (ATCC CRL-12203) were obtained from ATCC and propagated in DMEM supplemented as above. BS-C-1 cells (ATCC CCL-26) were purchased from ATCC and grown in minimum essential medium with Eagle’s balanced salts (EMEM) supplement as above.

The FAM111A-KO cell line was constructed using the CRISPR-Cas9 technology described previously^34^. Briefly, two different sgRNAs were designed to target human FAM111A (5’-AAAAATCCAGAAGACCAGACCA-3’, 5’-TGGGAAAATGTAATGACCACCT-3’) and were inserted into pSpCas9(BB)-2A-GFP, a gift from Feng Zhang (Addgene plasmid #48138). A549 cells were transfected with the above plasmids at 60% confluency and sorted by flow cytometry based on the expression of GFP at 48 h post-transfection. GFP-positive cells were serially diluted for clonal selection and successful inactivation of FAM111A was confirmed by Sanger sequencing and Western blotting analysis with a FAM111A antibody (Abcam).

The A549-ATG16L1-KD cell line stably expressing ATG16L1-shRNA was constructed with retroviral transduction. ATG16L1 shRNA (5’-GTCATCGACCTCCGGACAAAT-3’) was inserted into pLKO.1 puro (Addgene plasmid #8453) to generate pLKO.1-ATG16L1-shRNA. Retroviruses were packaged in 293T cells by co-transfecting the cells with 1ug pLKO.1-ATG16L1-shRNA, 1µg pMD2.G (Addgene plasmid #12259) and 1µg psPAX2 (Addgene plasmid #12260) in each well of a 6-well plate. Virus particles were collected 48 h post-transfection and filtered (0.22 μm) to remove cell debris. A549 cells were transduced with retroviruses at various dilutions in the presence of 5 μg/mL of polybrene for 48 h. Cells were passaged three times in a complete medium supplemented with 1 μg/mL of puromycin (Sigma-Aldrich). The A549-ATG16L1-KD and the A549-shControl cells were both maintained in the presence of 1 μg/mL puromycin and 500 μg/mL geneticin. The expression of ATG16L1 was determined by Western blotting using an anti-ATG16L1 antibody (Cell Signaling Technology).

### Antibodies

Rabbit antibody to VACV-WR strain and mouse antibody to VACV-I3 or VACV-E3 were described previously^35, 36^. Mouse anti-β-actin antibody (#AA128), mouse anti-histone H3 antibody (#AF0009), mouse anti-Flag antibody (#AF519), mouse anti-HA antibody (#AH158), HRP-labeled goat anti-mouse IgG (#A0216) and HRP-labeled goat anti-rabbit IgG (#A0208) were purchased from Beyotime Biotechnology; rabbit anti-FAM111A antibody (#ab184572), mouse anti-PCNA antibody (#ab29) and mouse anti-nuclear pore complex proteins antibody (#ab24609) were purchased from Abcam; rabbit anti-cGAS antibody (#79978), rabbit anti-STING antibody (#13647), rabbit anti-p-STING antibody (#19781), rabbit anti-LC3B antibody (#3868) and rabbit anti-ATG16L1 antibody (#8089) were purchased from Cell Signaling Technology; Goat anti-mouse IgG Alexa Fluor 488 (#A32723) and Goat anti-rabbit IgG Alexa Fluor 555 (#A32732) were purchased from Thermo Fisher Scientific.

#### Generation of recombinant vaccinia viruses

Recombinant viruses were constructed by homologous recombination using fluorescent reporter genes (mCherry or eGFP) for plaque selection. MVA+SPI-1 was previously described^34^. Briefly, a DNA cassette containing the C12L ORF regulated by the mH5 promoter followed by the GFP ORF regulated by the P11 promoter was generated and flanked by 5’ and 3’ sequences around the deletion III locus of MVA. CEF cells were infected with MVA at 3 PFU/cell and then transfected with the DNA cassette described above. Viruses were harvested 24 hpi and recombinant viruses were selected by selecting mCherry or eGFP positive clones and plaque-purified 3× times in CEF cells. Viral DNA was isolated from the clones with the DNeasy Blood & Tissue kit from Qiagen and the insertion of genes was verified by PCR amplification followed by Sanger sequencing. To generate MVA+SPI-1/C16, the C16L DNA segment along with the upstream 339 bp presumed to contain the natural promoter was copied by PCR from the genome of MVA-51.2 virus and inserted into a cassette that contained the mCherry ORF regulated by the P11 promoter with flanking sequences comprised of ORFs 069 and 070^18^. Homologous recombination was performed as above. All recombinant MVAs were propagated in CEF. VACV-ΔSPI-1 recombinant viruses were generated as follows. First, 500 bp up- and downstream of C12 ORF were cloned to the 5’ and 3’ end, respectively, of eGFP ORF regulated by the VACV p11 promoter. The resultant 1,720-bp PCR product was then transfected into BS-C-1 cells infected with VACV-WR at 3 PFU/cell for 1 hour. A recombinant virus lacking SPI-1 was isolated by selecting GFP-positive virus clones and clonally purified by repeated plaque isolation. The loss of the C12 gene was confirmed by PCR.

### Western Blotting analysis

Human A549 cells were washed once with ice-cold PBS and lysed in wells with 1× cell lysis buffer and 1× PMSF (Beyotime Biotechnology) for Western blotting analysis on ice. Cell lysates were sonicated for 1 min to reduce viscosity. Proteins were resolved on 4–20% FastPAGE precast gels (Tsingke Biotechnology) and transferred to nitrocellulose membranes in a Trans-Blot Turbo machine (Bio-Rad). Membranes were blocked with 5% nonfat milk dissolved in Tris-buffered saline (TBS) for 1 h at RT and then incubated with primary antibodies diluted 1× TBS with 0.1% Tween 20 (TBST) containing 5% nonfat milk overnight at 4 °C. The membranes were washed three times with 1× TBST and incubated with a secondary antibody conjugated with horseradish peroxidase diluted 1:5,000 in 1× TBST containing 5% nonfat milk for 1 hour at RT. ECL signals were detected with SuperSignal West Dura substrates (Thermo Fisher Scientific) and visualized by Image J.

### Immunoprecipitation

Human A549 cells were co-transfected with vectors expressing myc-tagged FAM111A and flag-tagged I3 or SPI-1 and total protein was harvested 24 hours post-transfection. Cells were first washed once with ice-cold PBS and lysed in wells with 150 µL 1× cell lysis buffer and 1× PMSF (Beyotime Biotechnology) in each well of a 6-well plate on ice. Protein lysates were centrifuged for 15 min at 15,000 × g at 4 C and then incubated with control magnetic beads, flag-trap magnetic beads, or myc-trap magnetic beads at 4 C for 3 hours with rotation. The beads were then washed 6× times with 1× lysis buffer and the bound proteins were eluted by boiling with 30 µL LDS sample buffer containing 0.05 M DTT for 15 min. The proteins were then resolved on 4–20% FastPAGE precast gels (Tsingke Biotechnology) and analyzed by Western blotting with antibodies described previously.

#### Determination of virus yield

Cell monolayers in 24-well plates were infected with viruses at a multiplicity of 0.03 PFU/cell in DMEM/F12 containing 2.5% FBS and 1% pen/strep for 2 hours. Cells were then washed twice with 1× phosphate-buffered saline (PBS) and incubated for 48 h. Viruses were harvested by scraping and followed by three freeze–thaw cycles. Duplicate DF-1 (for MVAs) or BS-C-1 (for VACV-WR and VACV-ΔSPI-1) monolayers were infected with serially diluted viruses for 2 h, and then the inoculum was aspirated and replaced with medium containing 2.5% FBS supplemented with 0.5% methylcellulose followed by 48h incubation. For quantification of MVA, DF-1 cells were fixed with methanol–acetone (1:1, vol/vol) and washed with distilled water, and incubated with anti-VACV-WR antibody diluted 1:1,000 in 3% FBS-PBS for 1 h. Cells were then washed twice with distilled water and incubated with a 1:3,000 dilution of protein A-HRP (Thermo Fisher Scientific). After 1 hour, protein A-HRP was aspirated and replaced with ethanol-saturated dianisidine, diluted 1:50 in PBS containing 0.3% H_2_O_2_. The dianisidine solution was removed after 10 min of incubation at room temperature, and cells were washed with distilled water. Plaques were counted manually. For quantification of VACV-WR and VACV-ΔSPI-1, cells were fixed with 1% crystal violet diluted in 20% EtOH, plaques were counted and average titers of triplicates were shown.

### Quantification of viral genomic DNA by quantitative PCR

Monolayers of A549 cells in 24-well plates were infected with indicated viruses at 3 PFU/cell for 1 hour in triplicates. Cells were washed twice with 1× PBS and total DNA was extracted at 6 hpi with DNeasy Blood/Tissue DNA mini kit (Qiagen) and quantitated by a nanodrop spectrophotometer (Thermo Fisher Scientific). Equal amounts (1µg) of DNA from each sample were then serially diluted and subjected to quantitative PCR with gene-specific primers for virus E11 and host GAPDH as the housekeeping control. The results of virus genomic DNA were normalized to that of GAPDH and relative DNA abundance was calculated.

#### Confocal microscopy

Human A549 cells and HeLa cells were cultured on glass coverslips and infected with viruses indicated in figures at 3 PFU/cell. Cells at various hpi were fixed with 4% paraformaldehyde (PFA) for 15 min, permeabilized with 0.5% Triton X-100 for 15 min, and blocked with 3% bovine serum albumin (BSA) in 1× PBS for 30 min at room temperature. Primary antibodies were diluted in 3% BSA-PBS and incubated with cells at 4 C overnight. Samples were then washed with 1× PBS, and incubated with secondary antibodies conjugated with Alexa Fluor 488 or Alexa Fluor 555. Nuclei and virus factories were stained with Hoechst and cells were washed multiple times before mounting on slides using ProLong Diamond Antifade reagent (Thermo Fisher Scientific). Images were taken on a Leica SP8 confocal microscope and images were processed with the Leica software (Leica Biosystems).

### Availability Statement

Materials and additional details regarding methods and protocols may be obtained by contacting the corresponding author via email.

## Acknowledgments

This work was funded by the National Key Research and Development Program (2021YFD1800700), the National Natural Science Foundation of China (32172822), and the Beijing STRS program (Z211100002121021) to Chen Peng.

## Contributions

**Conceived and designed the study:** Bernard Moss, Chen Peng, Junda Zhu

**Investigation:** Junda Zhu, Xintao Gao, Zihui Zhang, Shijie Xie, Shuning Ren, Yarui Li, Hua Li, Kang Niu, Shufang Fu, Yinü Li.

**Methodology:** Junda Zhu.

**Resources:** Chen Peng.

**Supervision:** Wenxue Wu, Chen Peng.

**Writing – original draft:** Chen Peng.

**Writing – review & editing:** Junda Zhu, Chen Peng, Bernard Moss

---

**Figure S1.**
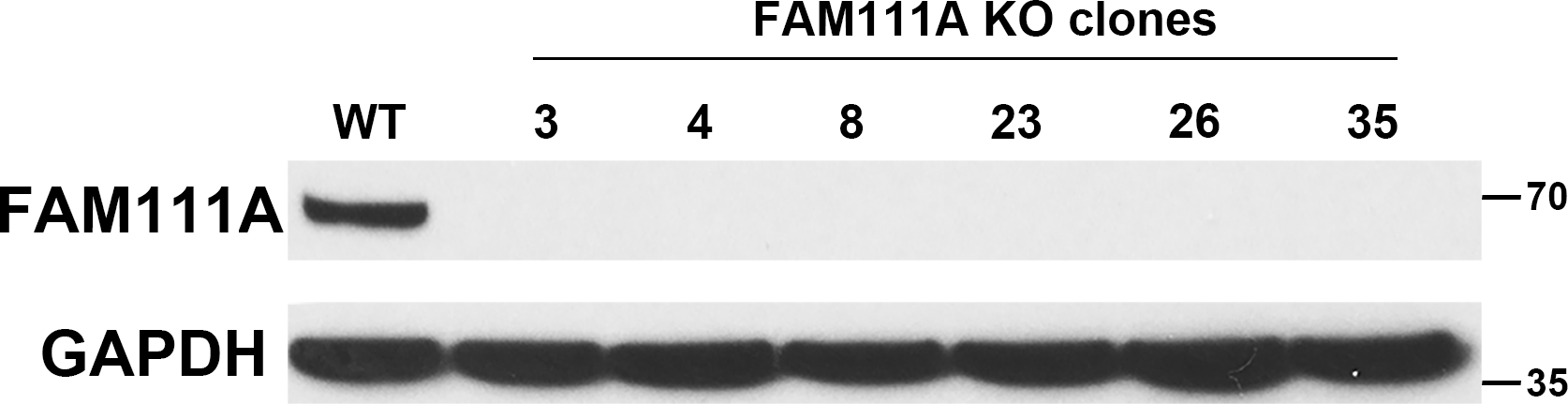
Cell clones of A549 FAM111A-KO by CRISPR-Cas9. A549^FAM111A-KO^ cells were constructed using CRISPR-Cas9 technology. A pool of A549 cells transfected with pSpCas9(BB)-2A-GFP containing sgRNA for FAM111A was diluted and single-cell clones were screened by Western blotting analysis using anti-FAM111A and anti-β-actin antibodies.

**Figure S2.**
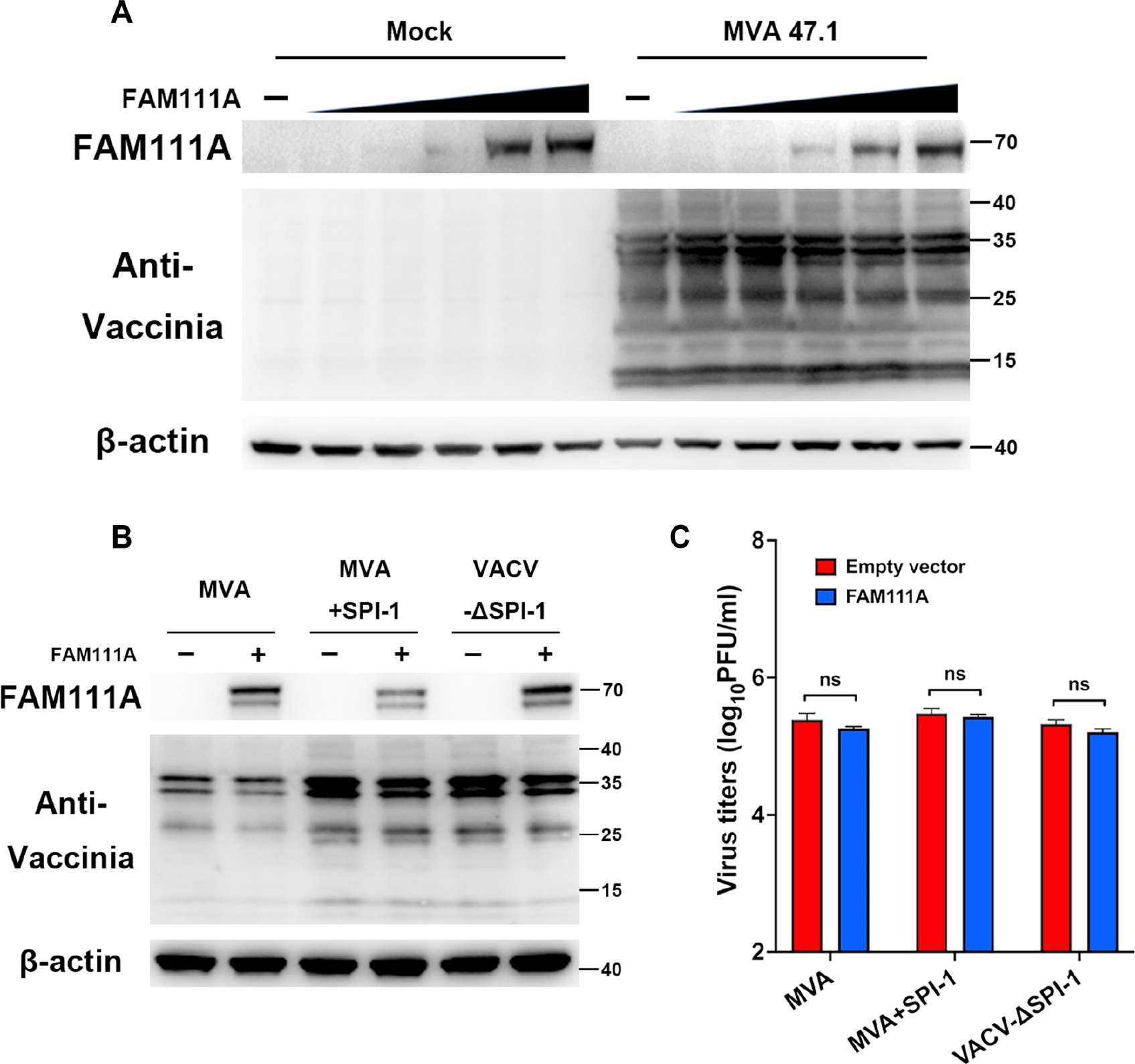
human-FAM111A does not expand the host range of MVA in BS-C-1 or DF-1 cells. **(A)** BS-C-1 cells were transfected with a vector expressing FAM111A at concentrations of 0.1, 0.2, 0.5, 1, and 2 μg/mL for 24 hours and then infected with MVA-47.1 at 0.3 PFU/cell. Cell lysates were analyzed by SDS-PAGE and Western blotting with anti- FAM111A, anti-VACV, or anti-β-actin antibodies at 24 hpi. **(B)** Avian DF-1 cells were transfected with a vector expressing FAM111A at concentrations of 1 μg/mL for 24 hours and then infected with MVA or MVA+SPI-1 at 0.3 PFU/cell. Cell lysates were analyzed by SDS-PAGE and Western blotting with anti-FAM111A, anti-VACV, or anti-β-actin antibodies at 4 hpi. **(C)** Chicken DF-1 cells were transfected with a vector expressing FAM111A at concentrations of 1 μg/mL for 24 hours and then infected with MVA, MVA+SPI-1, or VACV-ΔSPI-1 at 0.03 PFU/cell. Viruses were harvested at 24 hpi and virus titers were determined by plaque assay using DF-1 cells. Bars represent the mean values of virus titers. Data in A-C are representative of 3 independent experiments.

**Figure S3.**
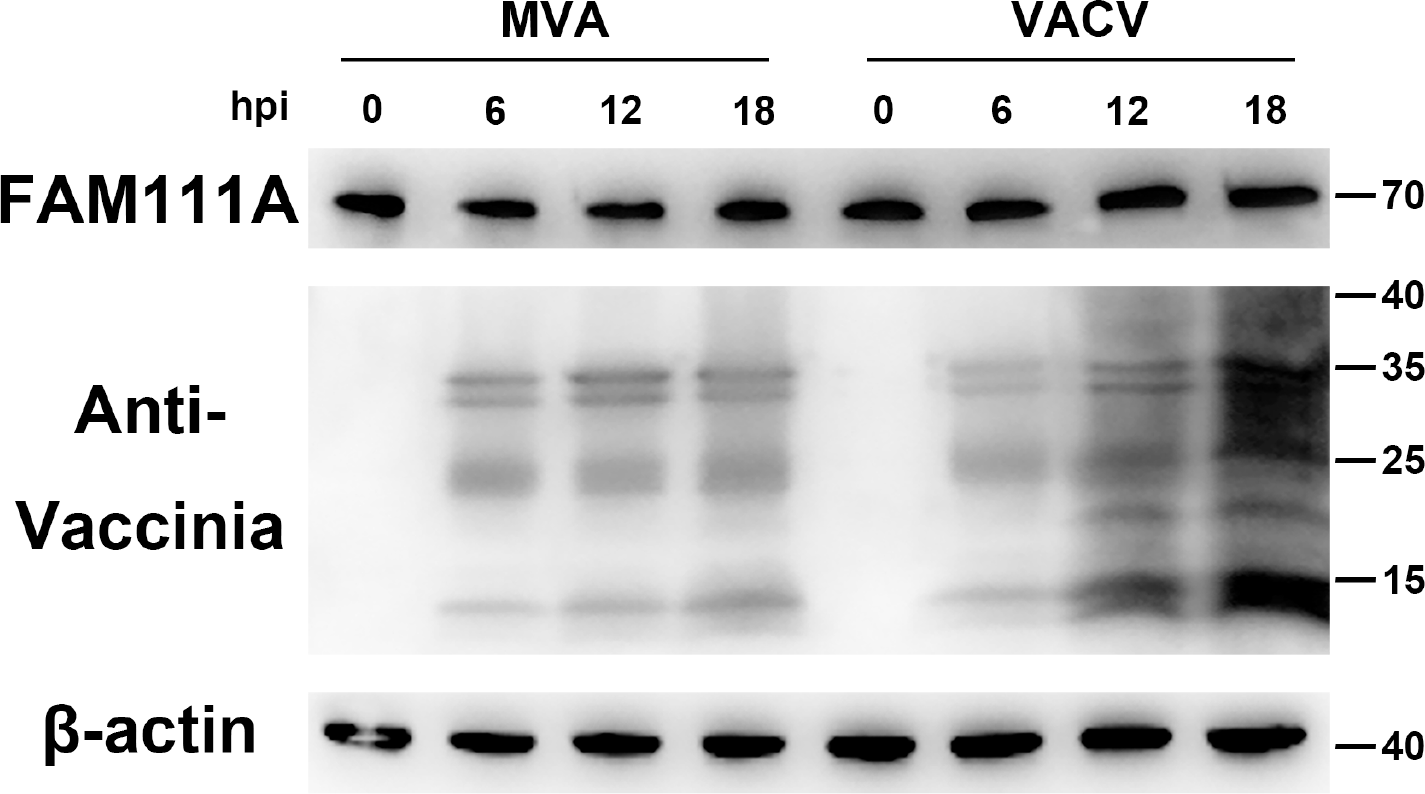
Expression levels of endogenous FAM111A after MVA and VACV-WR infection. Human A549 cells were infected with MVA or VACV-WR for 0, 6, 12, and 18 hours at 3 PFU/cell. Cell lysates were analyzed by SDS-PAGE and Western blotting with anti- FAM111A, anti-VACV, or anti-β-actin antibodies. Data are representative of 3 independent experiments.

**Figure S4.**
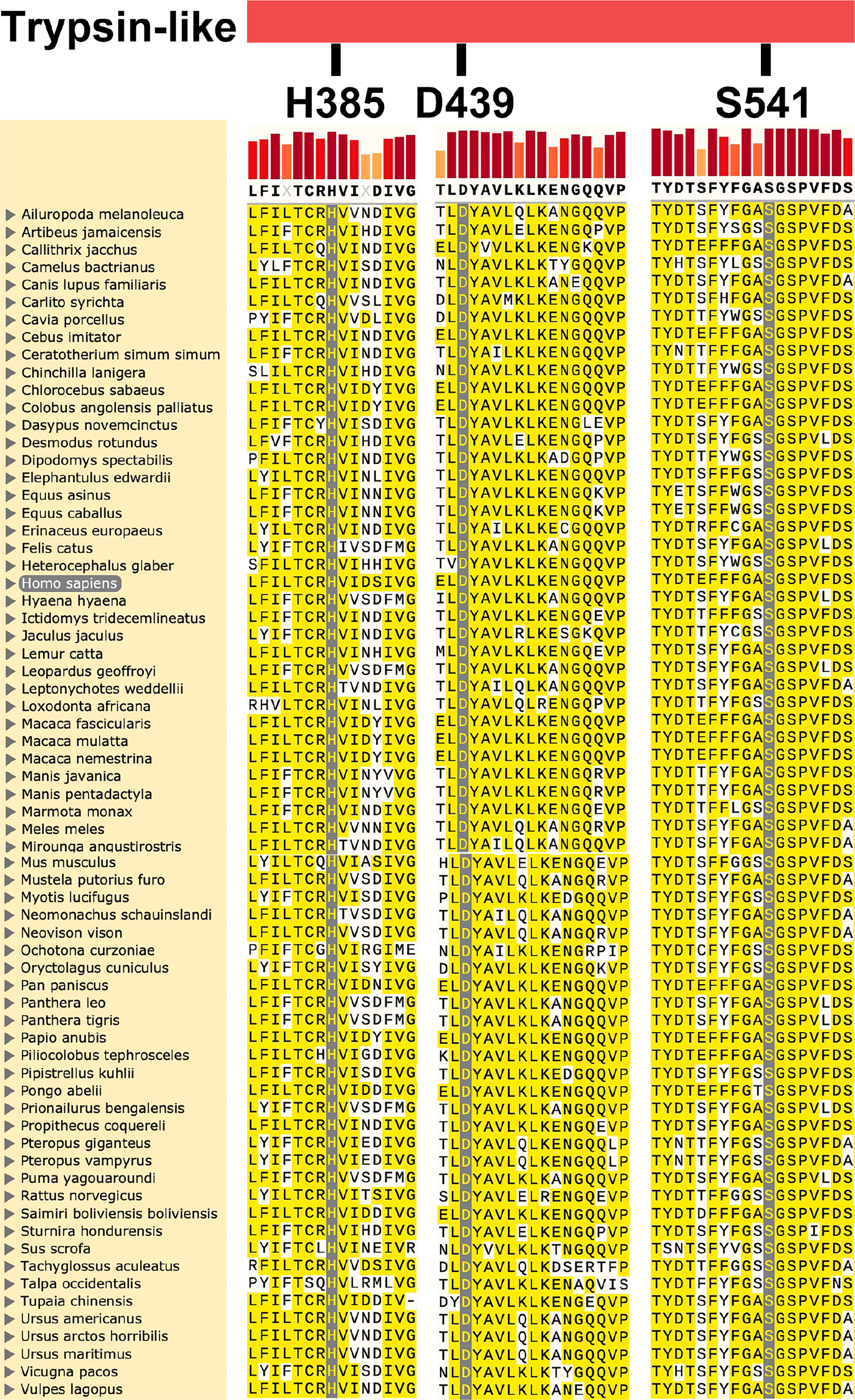
The serine protease catalytic triad at the C-terminus of FAM111A is highly conserved across mammals. The FAM111A amino acid sequences of 68 mammal species were downloaded from NCBI (https://www.ncbi.nlm.nih.gov/), and Snapgene was used for multiple sequence alignment for the C-terminal region containing amino acids 378 to 392, 437 to 453, and 531 to 548. Fully conserved residues are highlighted in grey and conserved residues in yellow.

**Figure S5.**
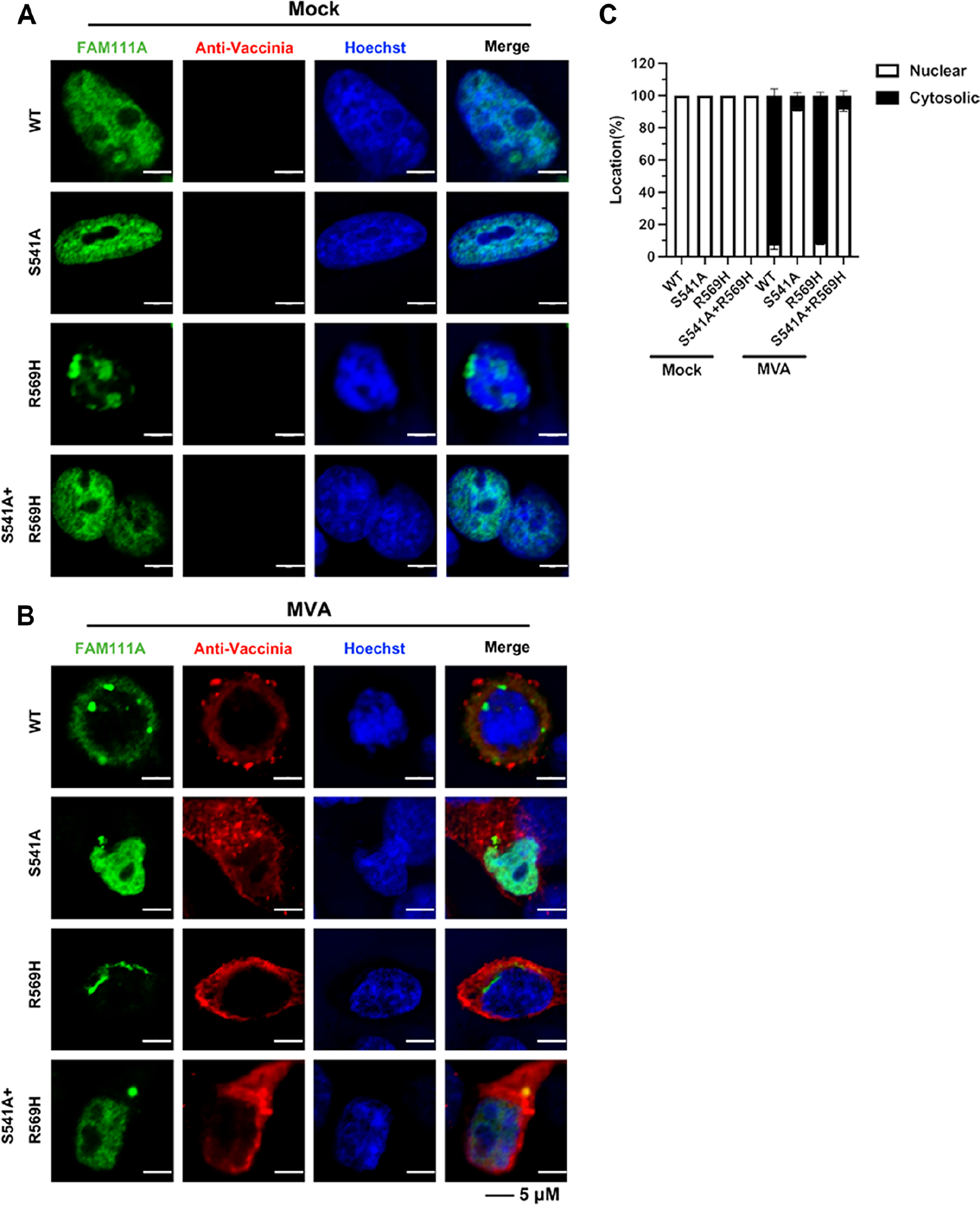
Virus infection triggers the protease-dependent relocalization of FAM111A in HeLa cells. **(A-C)** Human HeLa cells plated on coverslips were transfected with vectors expressing FAM111A^WT^, FAM111^AS541A^, FAM111A^R569H,^ or FAM111A^S541A+R569H^ and then mock infected or infected with MVA at 3 PFU/cell. At 12 hpi, cells were then fixed, permeabilized, blocked, and stained with primary antibodies to FAM111A and VACV followed by fluorescent conjugated secondary antibodies. Hoechst was used to stain DNA **(A and B)**. The confocal analyses were performed in triplicates and the localization of FAM111A was quantified in 100 randomly selected cells and bars represent mean values +/- SD **(C)**. Data in C represent the mean values ± SD (standard deviation) of 3 independent experiments. Data in A and B are representative of 3 independent experiments.

**Figure S6.**
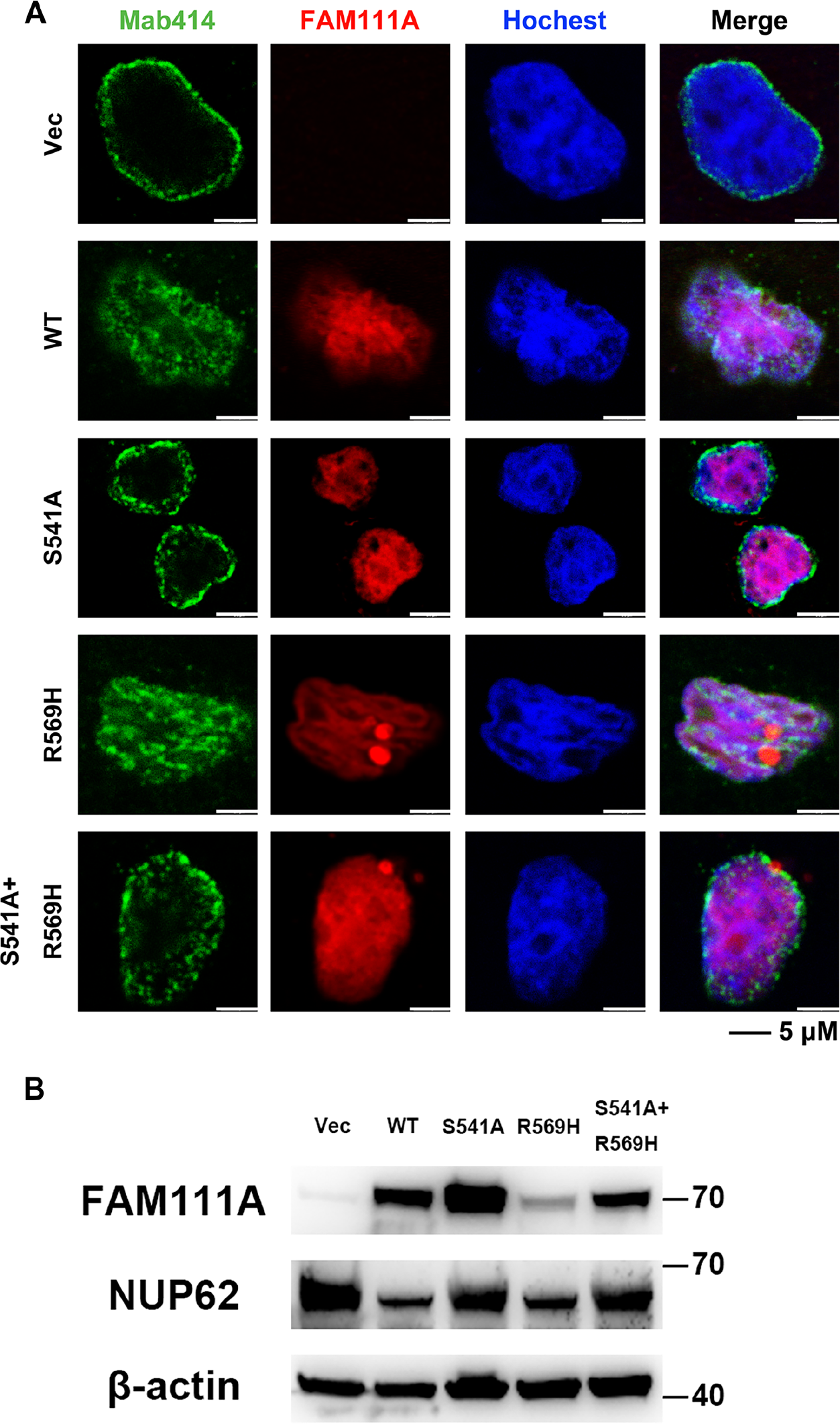
Protease-dependent disruption of the NPC upon overexpression of FAM111A. **(A)** A549 cells were transfected with vectors expressing FAM111A^WT^ and lab-generated mutants indicated in figure 2(A) for 24 hours. The cells were then fixed, permeabilized, blocked, and stained with primary antibodies to Mab414 and FAM111A followed by fluorescent conjugated secondary antibodies. Hoechst was used to stain DNA. Scale bar at bottom. **(B)** A549 cells were transfected with vectors expressing FAM111A^WT^ and indicated mutants for 24 hours. Cell lysates were analyzed by SDS-PAGE and Western blotting analysis using anti-FAM111A, Mab414 (anti-NUP62), and anti-β-actin antibodies. Data are representative of 3 independent experiments.

**Figure S7.**
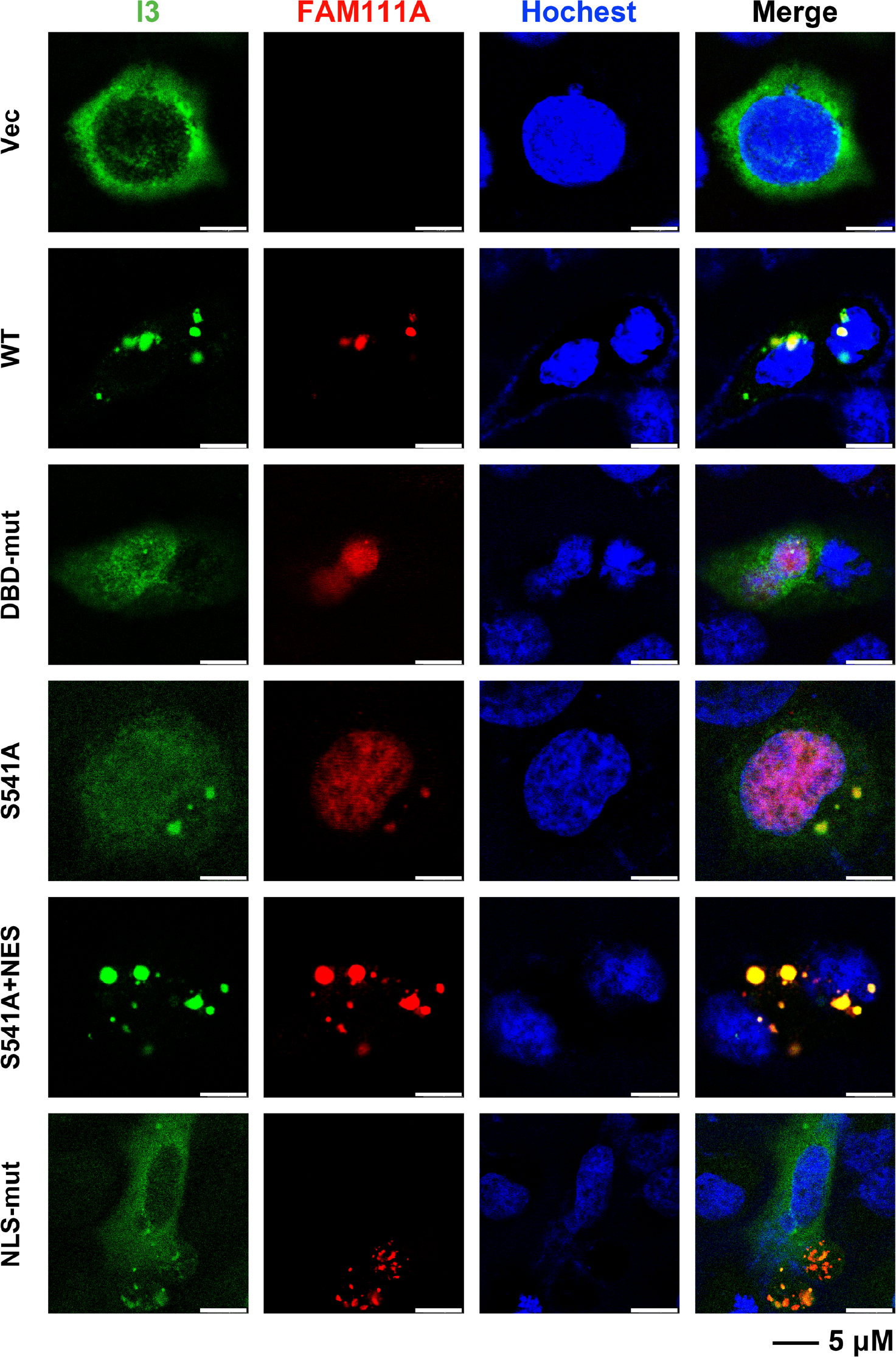
Subcellular localization of FAM111A and I3 in A549 cells. Human A549 cells were transfected with vectors expressing FAM111A^WT^ and indicated mutants for 24 hours. Then indicated cells were transfected with VACV-I3. After 24 hours, cells were then fixed, permeabilized, blocked, and stained with primary antibodies to I3 and FAM111A followed by fluorescent conjugated secondary antibodies. Hoechst was used to stain DNA. Scale bars are shown at bottom. Data are representative of 3 independent experiments.

**Figure S8.**
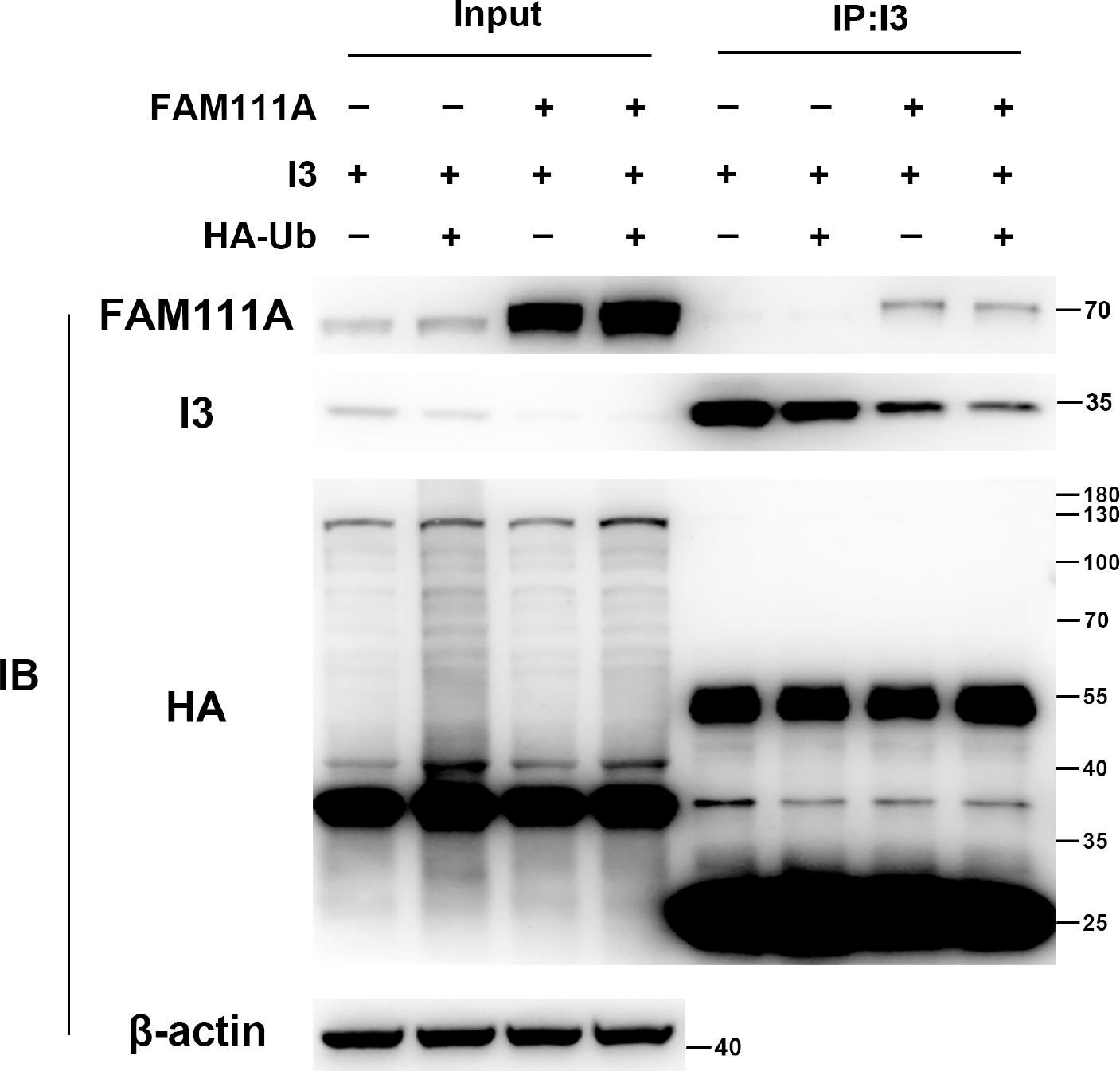
FAM111A does not degrade I3 through the ubiquitin-proteasome pathway. A549 cells were co-transfected with myc-tagged FAM111A, flag-tagged I3, and HA-tagged Ub for 24 hours. Cell lysates were incubated with control magnetic beads or flag-conjugated beads at 4°C for 18 hours. Beads were extensively washed and proteins were eluted with SDS-loading buffer and analyzed by SDS-PAGE and Western blotting analysis. The light chain or heavy chain antibody bands are at 25 kDa or 55 kDa, respectively. Data are representative of 3 independent experiments.

**Figure S9.**
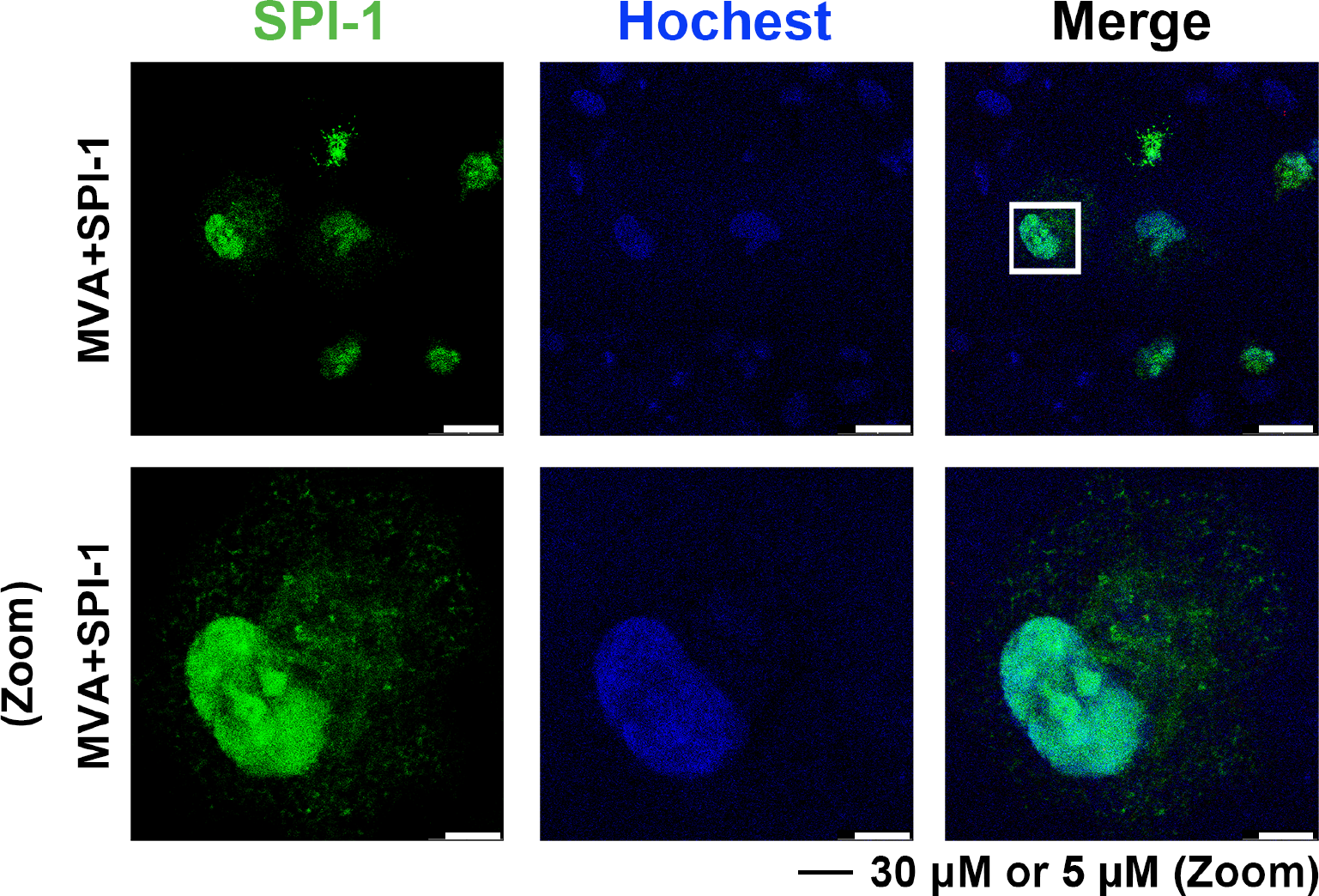
Subcellular localization of SPI-1 in A549 cells upon virus infection. Human A549 cells were infected with MVA+SPI-1 at 3 PFU/cell. After 24 hours, the cells were then fixed, permeabilized, blocked, and stained with primary antibodies to SPI-1 followed by fluorescent conjugated secondary antibodies. Hoechst was used to stain DNA. Scale bar at bottom. Data are representative of 3 independent experiments.

